# A lipocalin mediates unidirectional haem biomineralization in malaria parasites

**DOI:** 10.1101/2020.02.18.954289

**Authors:** Joachim M. Matz, Benjamin Drepper, Thorsten B. Blum, Eric van Genderen, Alana Burrell, Peer Martin, Thomas Stach, Lucy Collinson, Jan Pieter Abrahams, Kai Matuschewski, Michael J. Blackman

## Abstract

During blood stage development, malaria parasites are challenged with the detoxification of enormous amounts of haem released during the proteolytic catabolism of erythrocytic haemoglobin. They tackle this problem by sequestering haem into bioinert crystals known as haemozoin. The mechanisms underlying this biomineralization process remain enigmatic. Here, we demonstrate that both rodent and human malaria parasite species secrete and internalize a lipocalin-like protein, PV5, to control haem crystallization. Transcriptional deregulation of *PV5* in the rodent parasite *Plasmodium berghei* results in inordinate elongation of haemozoin crystals, while conditional *PV5* inactivation in the human malaria agent *Plasmodium falciparum* causes excessive multi-directional crystal branching. Although haemoglobin processing remains unaffected, PV5-deficient parasites generate less haemozoin. Electron diffraction analysis indicates that despite the distinct changes in crystal morphology neither the crystalline order nor unit cell of haemozoin are affected by impaired PV5 function. Deregulation of *PV5* expression renders *P. berghei* hypersensitive to the antimalarial drugs artesunate, chloroquine, and atovaquone, resulting in accelerated parasite clearance following drug treatment *in vivo*. Together, our findings demonstrate the *Plasmodium*-tailored role of a lipocalin family member in haemozoin formation and underscore the haem biomineralization pathway as an attractive target for therapeutic exploitation.

**SIGNIFICANCE:** During blood stage development, the malaria parasite replicates inside erythrocytes of the vertebrate host, where it engulfs and digests most of the available haemoglobin. This results in release of the oxygen-binding prosthetic group haem, which is highly toxic in its unbound form. The parasite crystallizes the haem into an insoluble pigment called haemozoin, a process that is vital for parasite survival and which is exploited in antimalarial therapy. We demonstrate that the parasite uses a protein called PV5 in haemozoin formation and that interfering with PV5 expression can increase the parasite’s sensitivity to antimalarial drugs during blood infection. An improved understanding of the mechanisms underlying haem sequestration will provide valuable insights for future drug development efforts.

## INTRODUCTION

The devastating pathology of malaria is caused by infection of red blood cells with unicellular *Plasmodium* parasites which reside within an intraerythrocytic parasitophorous vacuole (PV) (1). Throughout blood stage development, the parasite ingests and catabolizes up to 80% of the host cell cytoplasm, facilitating amino acid acquisition and making sufficient room for parasite growth (2, 3). Haemoglobin is incorporated through endocytosis and then degraded by an array of functionally redundant proteases, a process which occurs in acidified lysosome-like organelles with species-specific morphology (4). In the rodent-infective parasite species *Plasmodium berghei*, one or more food vacuoles (FVs) give rise to small digestive vesicles (DVs) which only fuse at the very end of intraerythrocytic development (5). By contrast, the most virulent agent of human malaria, *Plasmodium falciparum*, directs all endosomal traffic to a single large FV, where proteolysis occurs (6).

Here, haemoglobin digestion is accompanied by the release of high levels of the porphyrin co-factor haem from the globin chains. Haem can damage proteins and lipids through various mechanisms, including the formation of free radicals (7). The unique challenge of haem detoxification is met by the parasite’s capacity to sequester the released haem into a bioinert crystalline product called haemozoin (Hz), which accumulates in the FV or DVs. Haem is initially oxidised to yield haematin, which then dimerizes through the reciprocal coordination of iron and propionate moieties. This molecular unit then assembles into Hz crystals which typically take the form of triclinic high aspect ratio parallelograms (8–10). By the end of intraerythrocytic development, all the Hz crystals are contained within a central residual body, which is eventually released upon parasite egress from the host cell and which contributes to the inflammatory responses associated with acute malaria (11). The mechanism of Hz formation is highly debated. While several studies suggest a physicochemical and autocatalytic crystallization process (12–15), there have been reports of parasite proteins (16–18) and lipids (19–21) promoting Hz assembly *in vitro*.

The parasite’s dependency on haem detoxification has long been exploited in antimalarial therapy with outstanding success. Aminoquinolines inhibit Hz formation *via* direct physical interactions with haematin and the crystal surface, eventually leading to the build-up of cytotoxic free haem (22–25). The aminoquinoline chloroquine was the front-line medication against malaria from the 1950s onward until the emergence of wide-spread drug resistance restricted its utility (26). Nonetheless, to this day, chloroquine remains among the most effective antimalarial drugs ever developed, highlighting the outstanding importance of haem sequestration for *Plasmodium* survival. It is thus crucial for future drug development efforts to gain a better understanding of the mechanisms underlying this unique biomineralization event.

In this report, we demonstrate that a parasite-encoded lipocalin called PV5 is a central regulator of *Hz* formation.

## RESULTS

### Malaria parasites encode a lipocalin-like protein, PV5

Employing a genome-wide *in silico* down-scaling approach, we previously identified an essential *P. berghei* PV protein, *Pb*PV5 (PBANKA_0826700), which has orthologues in all other *Plasmodium* species, including *P. falciparum* (PF3D7_0925900) (27). Inspection of the PV5 amino acid sequence revealed a striking similarity to members of the functionally diverse lipocalin family, barrel-shaped proteins capable of binding various hydrophobic ligands and/or protein interaction partners (28). The signature lipocalin fold comprises a short amino-terminal helix followed by eight consecutive barrel-forming β-strands, another α-helix and one more β-strand (Fig. 1*A*) (29). In addition, PV5 harbours two preceding amino-terminal β-strands specific to *Plasmodium*, as well as a signal peptide. Multiple sequence alignments with lipocalins from phylogenetically distant organisms showed the presence of a highly conserved glycine and two aromatic amino acids within the structurally conserved region 1 (SCR1) of PV5, a hallmark of the extended calycin superfamily (Fig. 1*A*) (29). Among several structural homologues, the bacterial outer membrane lipocalin Blc from *Escherichia coli* was predicted to share the highest similarity with PV5. Homology modelling guided by the known *E. coli* Blc structure suggests that PV5 shares the overall architecture of the lipocalin family including the characteristic β-barrel (Fig. 1*B*). Together, the sequence signatures and predicted structural features support membership of *Plasmodium* PV5 in the calycin protein superfamily.

**Fig. 1.**
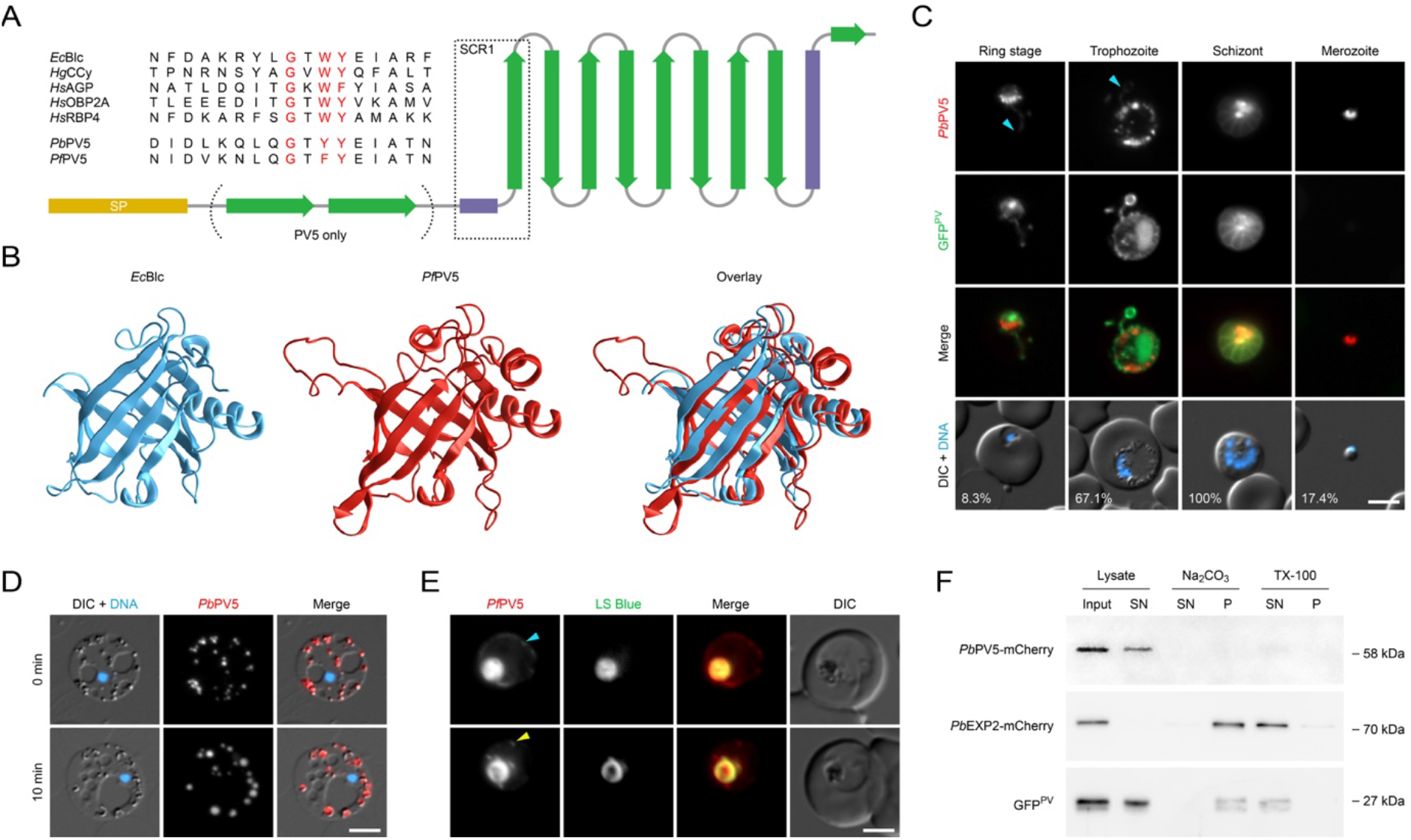
The *Plasmodium* lipocalin PV5 is trafficked to the parasite’s digestive compartments. (*A*) PV5 is a lipocalin family member. Secondary structure of *Plasmodium* PV5. Yellow, signal peptide (SP); green, ß-strands; purple, α-helices. Note the two amino-terminal ß-strands specific to PV5. Alignments of the structurally conserved region 1 (SCR1) from different lipocalin family members and PV5 are shown in the upper left corner. Signature residues are highlighted in red. *Ec*Blc, *Escherichia coli* bacterial lipocalin; *Hg*CCy; *Homarus gammarus* (European lobster) crustacyanin; *Hs*AGP, *Homo sapiens* α_1_-acid glycoprotein; *Hs*OBP2A, *H. sapiens* odorant-binding protein 2A; *Hs*RBP4, *H. sapiens* retinol-binding protein 4; *Pb*/*Pf*PV5, PV5 from *P. berghei* and *P. falciparum*. (*B*) Structure homology modelling predicts a lipocalin fold for *Pf*PV5. Shown are the experimentally validated structure of *Ec*Blc (blue, left, residues 27-175, PDB ID: 3MBT), the derived model of *Pf*PV5 (red, centre, residues 35-214) using *Ec*Blc as a homology template, and an overlay (right). Modelling was performed with SWISS-MODEL and supported by I-TASSER. Amino acid sequence identity is 20%, similarity calculated from BLOSUM62 substitution matrix is 0.3. (*C*) Dual protein localization of PV5 to extensions of the PV and to intraparasitic structures in *P. berghei*. Transgenic parasites expressing the PV marker GFP^PV^ and the endogenous Pb*PV5* gene fused to mCherry-3xMyc were imaged live (27). Shown are the mCherry (red, first row) and GFP channels (green, second row), a merge of both signals (third row) and a merge of differential interference contrast images (DIC) and Hoechst 33342 nuclear stain (blue, fourth row). Cyan arrowheads, *Pb*PV5 in PV tubules. Numbers represent normalized mCherry intensity values obtained by quantitative live fluorescence microscopy. N=44 parasites. (*D*) Intraparasitic PV5 localizes to Hz-containing DVs in *P. berghei*. Parasites were incubated in 1-2 μl under a large coverslip (22 x 40 mm) for several minutes, leading to lysis of the host erythrocyte and the PV, and to mechanical expansion of the parasite (top). Shown are a merge of DIC and Hoechst 33342 nuclear stain (blue, first row), the signal of tagged *Pb*PV5 (red, second row), as well as a merge of all three channels (third row). Swelling of *Pb*PV5-containing DVs was observed ten minutes later (bottom). Note the even distribution of *Pb*PV5 throughout the swollen DVs. (*E*) PV5 localizes to the central FV, intraparasitic vesicles and to the PV in *P. falciparum*. Transgenic parasites expressing the endogenous Pf*PV5* gene fused to mCherry were imaged live in the presence of Lysosensor Blue DND-167 (LS Blue). Shown are the signals of mCherry (red, first row), LS Blue (green, second row), a merge of both signals (third row), and DIC images (fourth row). Cyan arrowhead, *Pf*PV5 in PV (top). Yellow arrowhead, *Pf*PV5 in small intraparasitic vesicles (bottom). Bars, 5 μm. (*F*) PV5 is a soluble protein. Subcellular fractionation was performed using the *Pb*PV5-tagged *P. berghei* line, which also expresses the soluble marker GFP^PV^, and a *P. berghei* line expressing the transmembrane protein *Pb*EXP2 fused to mCherry-3xMyc (64). Cell lysates were centrifuged and resultant membrane pellets were subjected to solubilization with Na_2_CO_3_ and Triton X-100 (TX-100). Input, supernatant (SN) and pellet fractions (P) were analysed by Western blot using anti-mCherry and anti-GFP primary antibodies.

### PV5 is trafficked to the parasite’s digestive compartments

We first investigated the spatiotemporal expression of *Pb*PV5 during asexual blood stage development. Live fluorescence microscopy of transgenic *P. berghei* parasites expressing mCherry-3xMyc-tagged *Pb*PV5 confirmed that the protein localizes to tubular PV extensions during ring and trophozoite stages and surrounds individual merozoites in segmented schizonts (Fig. 1*C*) (27). In addition, a substantial fraction of the protein was restricted to the parasite cytoplasm. This was particularly prominent in schizonts, where intraparasitic *Pb*PV5 appeared to localize to the Hz-containing residual body (Fig. 1*C*). In merozoites, mCherry fluorescence was concentrated in a punctate intraparasitic region, perhaps signifying storage of *Pb*PV5 in the dense granules, as has been demonstrated for several other important PV proteins (30–32), but this fraction was minimal as compared to the protein contained in the residual body. Quantification of the mCherry-fluorescence intensity in live parasites indicated that *Pb*PV5 is much more abundant in mature parasite stages than in rings and merozoites, suggesting substantial levels of *de novo* synthesis throughout parasite maturation (Fig 1*C*).

In trophozoites, the intraparasitic fraction of *Pb*PV5 was associated with spherical structures at the parasite periphery (Fig. 1*C*) and microscopic examination of mechanically expanded free parasites revealed that these were the Hz-containing DVs (Fig. 1*D*.) To test whether this localization is conserved across different *Plasmodium* species, we generated similar transgenic *P. falciparum* parasites expressing mCherry-tagged *Pf*PV5. Here, a small fraction of the fusion protein consistently localized to the PV, whereas the majority of the fluorescent signal overlapped with the Hz crystals and with the signal of the acidotropic dye Lysosensor Blue DND-167, which accumulates in the acidic FV (Fig. 1*E*). In addition, we frequently observed smaller *Pf*PV5-positive foci in the parasite cytoplasm, most likely reflecting early endosomal compartments (Fig. 1*E*). Subcellular fractionation of tagged *P. berghei* parasites revealed that *Pb*PV5 is freely soluble (Fig. 1*F*). These finding are in good agreement with the detection of *Pf*PV5 in the FV proteome of *P. falciparum* (33). Together, our observations suggest that in both *Plasmodium* species PV5 is first secreted into the PV and then internalized through endocytosis of host cell cytoplasm to accumulate in the matrix of the parasite’s digestive compartments.

### Transcriptional deregulation of Pb*PV5* impairs asexual parasite propagation *in vivo*

Our previous attempts to disrupt the genomic Pb*PV5* locus resulted only in atypical integration of the targeting construct without perturbing the endogenous gene, which is indicative of essential functions during the asexual blood stage cycle *in vivo* (27). As an alternative genetic strategy to analyse Pb*PV5* function, we sought to deregulate Pb*PV5* expression by employing a promoter swap approach (Fig. 2*A*). Towards this aim, we generated parasites expressing the endogenous Pb*PV5* gene from the promoters of *Plasmodium* translocon of exported proteins 88 *(PTEX88)* or heat shock protein 101 *(HSP101)*, respectively (Fig. S1 *A* and *B*). Quantitative real-time PCR analysis of the mutants indicated that the knock-down efficiency was ~60% in asynchronous blood stages (Fig. S1*C*). Impaired growth prevented quantification of knock-down levels in synchronized *ex vivo* early blood stages. Strikingly, in the schizont stage, mutants exhibited significantly elevated Pb*PV5* transcript levels, corresponding to ~3.4 *(pv5::5’ptex88)* and ~5.6 times (*pv5::5’hsp101)* more than in WT schizonts (Fig. S1*C*). Therefore, using the promoter swap strategy, we succeeded in deregulating the physiological expression of Pb*PV5* throughout the asexual replication cycle.

**Fig. 2.**
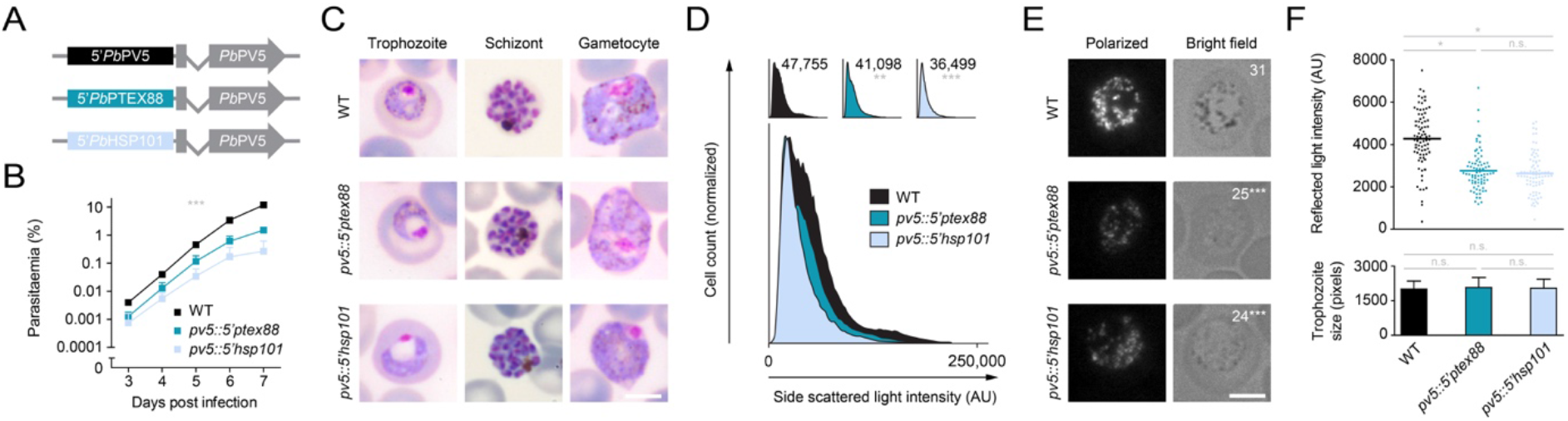
Deregulated expression of *PV5* impacts haemozoin formation in *Plasmodium berghei*. *(A)* Schematic representation of the genotypes of WT (top) and transgenic *pv5::5’ptex88* (middle) and *pv5::5’hsp101* parasites (bottom). In the mutants, the endogenous Pb*PV5* promoter (black) was exchanged for the promoter of *PbPTEX88* (dark blue) or *PbHSP101* (light blue), respectively. (*B*) Reduced parasite proliferation upon *Pb*PV5 promoter swapping. Asexual blood stage development was analysed using the intravital competition assay (55). Average parasite multiplication rates were 11.4 (WT), 9.1 *(pv5::5’ptex88)* and 6.2 *(pv5::5’hsp101*.) Shown are mean values +/-SD. ***, P<0.001; Two-way ANOVA. N=3 independent infections. (*C*) Morphology of trophozoite, schizont and gametocyte stages in the WT, *pv5::5’ptex88* and *pv5::5’hsp101* lines as observed by Giemsa staining. Note the lack of prominent dark pigment granules in mutant trophozoites and gametocytes as well as the dilation of the FV in mutant trophozoites. Bar, 5 μm. (*D*)Pb*PV5* promoter swap mutants are less granular. Infected blood was subjected to flow cytometry and the intensity of the side scattered light was determined. Shown are individual histograms including the mean side scatter intensity values (top) as well as a merge of WT, *pv5::5’ptex88* and *pv5::5’hsp101* histograms (bottom). Significances are shown for the comparison of the mutants with WT. n.s., non-significant; **, P<0.01; ***, P<0.001; One-way ANOVA and Tukey’s multiple comparison test. N=6 independent infections. *(E, F) Pb*PV5 is required for efficient Hz formation. (*E*) Trophozoites were visualized by polarization microscopy (left) and bright field imaging (right). Numbers indicate the mean quantity of bright puncta in polarization images. Significances are shown for the comparison of the mutants with WT. Bar, 5 μm. (*F*) Quantitative polarization microscopy. Depicted are individual and mean intensity values of reflected polarized light in methanol-fixed WT, *pv5::5’ptex88* and *pv5::5’hsp101* trophozoites (bars, upper graph). Only trophozoites of similar size were analysed (lower graph). Depicted are mean values +/-SD. n.s., non-significant; *, P<0.05; ***, P<0.001; One-way ANOVA and Tukey’s multiple comparison test. N=80 trophozoites from 4 independent infections.

To investigate whether altered Pb*PV5* transcription results in reduced parasite fitness, we examined asexual propagation of the mutants *in vivo*. Growth of the promoter swap mutants was significantly impaired, with the *pv5::5’ptex88* parasites growing at 80% and *pv5::5’hsp101* parasites at only 54% of the WT growth rate (Fig. 2*B*). These results underline the importance of correct Pb*PV5* expression during asexual replication of the parasite *in vivo*.

### Pb*PV5* mutants form less haemozoin

Inspection of Giemsa-stained thin blood films revealed striking morphological differences between WT parasites and the promoter swap mutants. During the trophozoite stage, the FV of the mutants was significantly swollen, visible as a large translucent area within the parasite cytoplasm close to the nucleus (Fig. 2*C*). Microscopic quantification revealed this area to be 1.5-fold (*pv5::5’ptex88*) or 1.8-fold (*pv5::5’hsp101*) larger than in WT parasites (Fig. S2*A*). Vacuolar swelling was transient, as mature schizonts did not exhibit comparable abnormalities (Fig. 2*C*).

Another striking phenomenon was the low visibility of dark granular material in mutant trophozoites, when compared to WT (Fig. 2*C*). This lack of granularity was most noticeable in the *pv5::5’hsp101* mutant and became particularly apparent during the gametocyte stage, where pigment granules are usually very prominent. To validate this finding, we subjected mixed blood stage parasites to flow cytometry and measured the intensity of the side scattered light, a commonly used proxy for cellular granularity (Fig. 2*D*). In agreement with our microscopic analysis, the Pb*PV5* mutants displayed reduced side scattering and the phenotype was again more severe in the *pv5::5’hsp101* mutant.

Because we suspected a Hz formation defect in the mutants, we fixed intraerythrocytic parasites with methanol and subjected them to polarization microscopy, which exploits the birefringent properties of Hz to specifically visualize the crystals. Using this method, we detected a weaker signal for the mutants, which correlated with reduced visibility of dark pigment in brightfield (Fig. 2*E*). Enumeration of individual bright entities suggested the presence of fewer Hz-containing structures within the mutants (~80% of WT) (Fig. 2*E*). Quantification of the polarized light intensity also indicated that *pv5::5’ptex88* parasites form only 64% and the *pv5::5’hsp101* 61 % of the Hz generated in WT parasites (Fig. 2*F*.) Together, these observations show that perturbation of Pb*PV5* expression results in reduced haem biomineralization *in vivo*.

### Protracted haemozoin extension upon deregulation of Pb*PV5* expression

We next aimed to examine how lower levels of Hz correlate with crystal size. To do this, we isolated Hz from mixed blood stage parasites and examined the material by scanning electron microscopy (SEM). This confirmed the characteristic high aspect ratio parallelogram morphology of WT Hz (Fig. 3 *A* and *B*). Strikingly, this was not the case for crystals isolated from the promoter swap mutants. Hz from both transgenic parasite strains exhibited highly irregular shapes and rough edges and showed only few of the distinctive Hz crystal vertices (Fig. 3*B*). Crystals from *pv5::5’ptex88* parasites most often had a pointed and canine tooth-like appearance, extending from a single straight crystal face. These abnormalities were even more pronounced in the *pv5::5’hsp101* mutant, where in most cases there was a region of normal crystal morphology with 2 or 3 straight edges, corresponding to the {010}, {011} and {001} faces. From this regular parent crystal emerged an enormous outgrowth which usually surpassed the dimensions of the parent crystal (Fig. 3*B*). This outgrowth consistently grew at an obtuse angle of ~105° in relation to the dominant *c* axis of the parent crystal, although accurate determination of the angle was complicated by the slightly bent and irregular shape of the outgrowth, which might be attributed to the space restrictions encountered in the *P. berghei* DVs. The outgrowth’s angle was not reflected in the physiological morphology of Hz (Fig. 3 *A* and *B*) and at least two faces of the regular parent crystal appeared to be involved. Indeed, in most cases, the outgrowth emerged from sites where the {010} and {011} faces meet and always grew along a plane corresponding to one of the original crystal faces (Fig. 3*B*). Other crystal formations were also observed, albeit at lower frequency, including some with multiple crystal branches and some with very rough surfaces (Fig. S2*B*). We observed similar crystal abnormalities *in situ* by transmission electron microscopy (TEM) of purified schizonts (Fig. 3*C*).

**Fig. 3.**
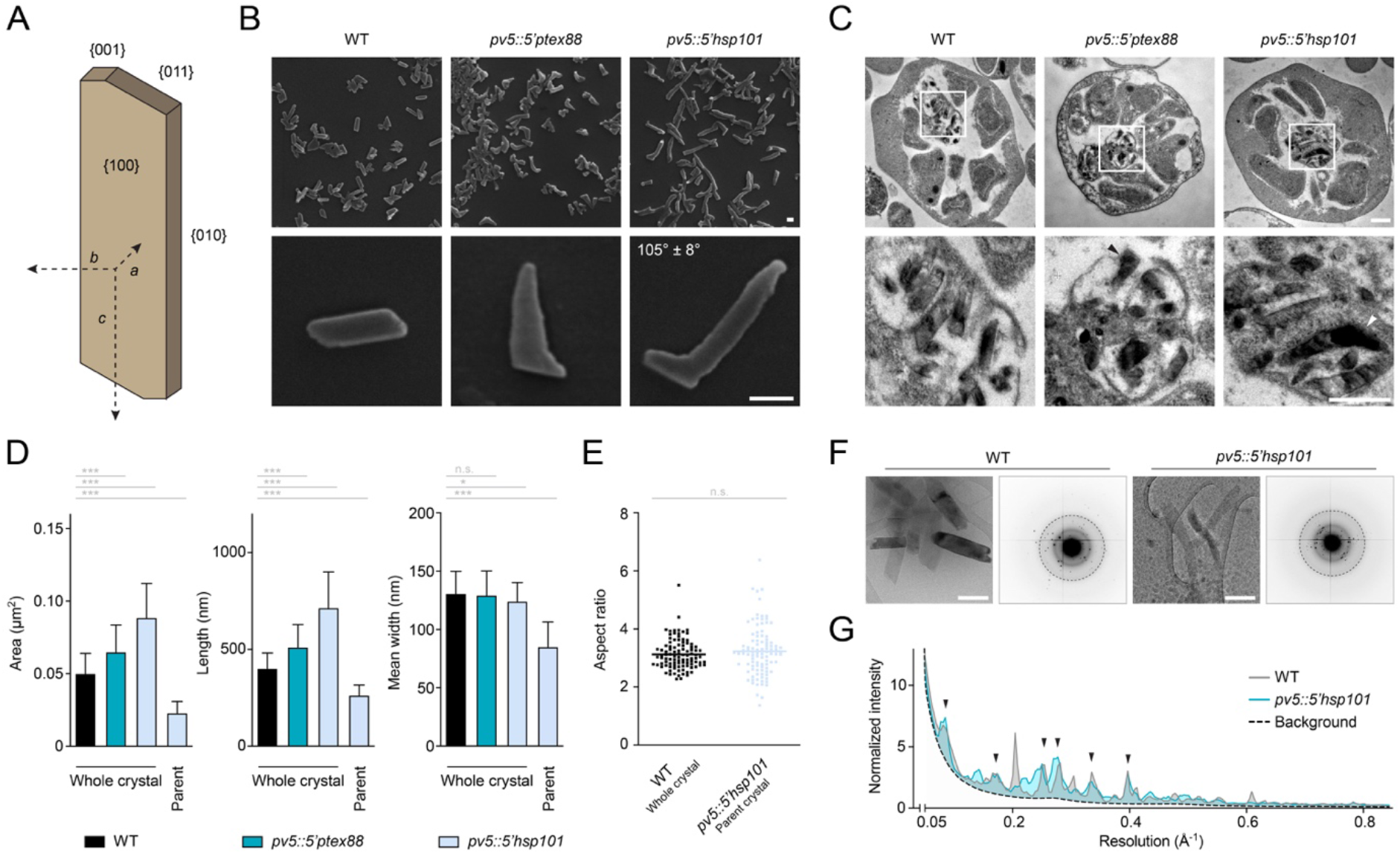
PV5 determines haemozoin morphology in *Plasmodium berghei*. (A) Hz crystal architecture. In *Plasmodium* WT parasites, Hz assembles as triclinic high aspect ratio parallelograms (63). Characteristic crystal axes and faces are indicated. (*B*) Scanning electron micrographs of Hz purified from WT, *pv5::5’ptex88* and *pv5::5’hsp101* mixed blood stage parasites. The angle between the regularly shaped *pv5::5’hsp101* parent crystal and the outgrowth is indicated. N=130 crystals. Bars, 100 nm. (*C*) Crystal morphology *in situ*. Shown are TEM images of WT, *pv5::5’ptex88* and *pv5::5’hsp101* schizonts (top) as well as their residual body at higher magnification (bottom). Abnormal crystal shapes in the mutants are indicated by arrowheads. Bars, 500 nm. *(D)* Hyperactive Hz growth is unidirectional. Shown are the dimensions of whole individual Hz crystals extracted from WT, *pv5::5’ptex88* and *pv5::5’hsp101* parasites including the area exposed to the electron beam (left) as well as the length (centre) and mean width of the crystals (right). The theoretical dimensions of the *pv5::5’hsp101* parent crystal were interpolated and are depicted as well. Shown are mean values +/-SD. n.s., non-significant; *, P<0.05; ***, One-way ANOVA and Tukey’s multiple comparison test. N=100 crystals. (*E*) Normal aspect ratio of *pv5::5’hsp101* parent crystals. Shown are the aspect ratios of whole Hz crystals from WT parasites and of the parent crystal from *pv5::5’hsp101*-generated Hz. Depicted are individual and mean values (bars). n.s., non-significant; Student’s *t*-test. N=100 crystals. *(F, G)* Unaltered crystalline order in Hz of *pv5::5’hsp101* parasites. (*F*) Depicted are TEM images (left) of Hz purified from WT and *pv5::5’hsp101* parasites as well as their corresponding electron diffraction patterns (right) showing comparable resolution of the Bragg peaks. Dashed circles demark a resolution of 0.5 Å^-1^. Bars, 200 nm. (*G*) Plot of the radial maximum diffracted intensity as a function of resolution. Data were normalized to the average median intensity at 0.18 Å^-1^, in order to correct for differences in diffracted volume. Arrowheads denote overlapping peaks. N=10 (WT) and 18 *(pv5::5’hsp101)* diffraction data sets.

Despite the lower overall levels of Hz formed and the abnormal crystal morphology, we found that the mutants formed larger Hz crystals as indicated by the area exposed to the SEM electron beam (Fig. 3*D*). The *pv5::5’hsp101* mutant formed the largest crystals, which were ~180% of WT size, while the *pv5::5’ptex88* crystals were at 130%. Importantly, the mutant Hz crystals displayed greater dimensions only in length but not in width owing to the unidirectional expansion of the outgrowth (Fig. 3*D*). Examination of the parent crystals from *pv5::5’hsp101* parasites showed that these were roughly half the size of whole WT crystals (Fig. 3*D*). The aspect ratios of WT crystals and the *pv5::5’hsp101* parent crystals were identical, together indicating that a period of normal crystallisation during earlier *pv5::5’hsp101* parasite development is followed by irregular crystal extension later on (Fig. 3*E*). Inspection of Hz from an unrelated slow-growing mutant (35) and from chloroquine-treated WT parasites ruled out a secondary effect of reduced parasite fitness or mortality on Hz morphology (Fig. S2 *C* and *D*).

To determine whether the crystalline order was affected by deregulation of Pb*PV5*, we obtained electron diffraction patterns from WT- and *pv5::5’hsp101*-derived Hz (Fig. 3*E*) and analysed the maximum diffracted intensities in concentric bins as a function of resolution. There was no difference in the drop-off of diffracted intensity between WT and mutant and the peak positions of the maxima corresponded (Fig. 3*G*). Differences in the magnitude of individual peaks can be attributed to preferential orientation, especially of the *pv5::5’hsp101*-derived crystals, which most often come to lie at their {100} faces. Together, these data indicate no differences in crystalline order or unit cell upon functional impairment of Pb*PV5* and we conclude that altered crystal morphology is not caused by alternative nucleation into a different haematin polymorph. Thus, deregulation of Pb*PV5* leads to the formation of ordered elongated Hz crystals with a highly variable and abnormal habit.

### Loss of PV5 causes haemozoin branching in *P. falciparum*

To investigate the consequences of *PV5* disruption, we generated a conditional *PV5* knockout line of the human pathogen *P. falciparum*, allowing rapamycin-induced DiCre-mediated excision of the Pf*PV5* gene (Fig. 4*A* and Fig. S3*A*). Correct modification of the locus was indicated by diagnostic PCR and by the successful tagging of *Pf*PV5 with a 3xHA tag, as demonstrated by Western blot and immunofluorescence analysis (Fig. 4 *B* and *C*, Fig. S3 *B*-*D*). 3xHA-tagged *Pf*PV5 localized to the PV and to intraparasitic vesicular structures (Fig. S3*C* and *D*). Surprisingly, no signal was observed within the central FV, in marked contrast to the mCherry fusion protein (Fig. 1*E*) and despite previous evidence from FV proteomics (33). Most likely, the 3xHA-tag is obscured or processed in the FV.

**Fig. 4.**
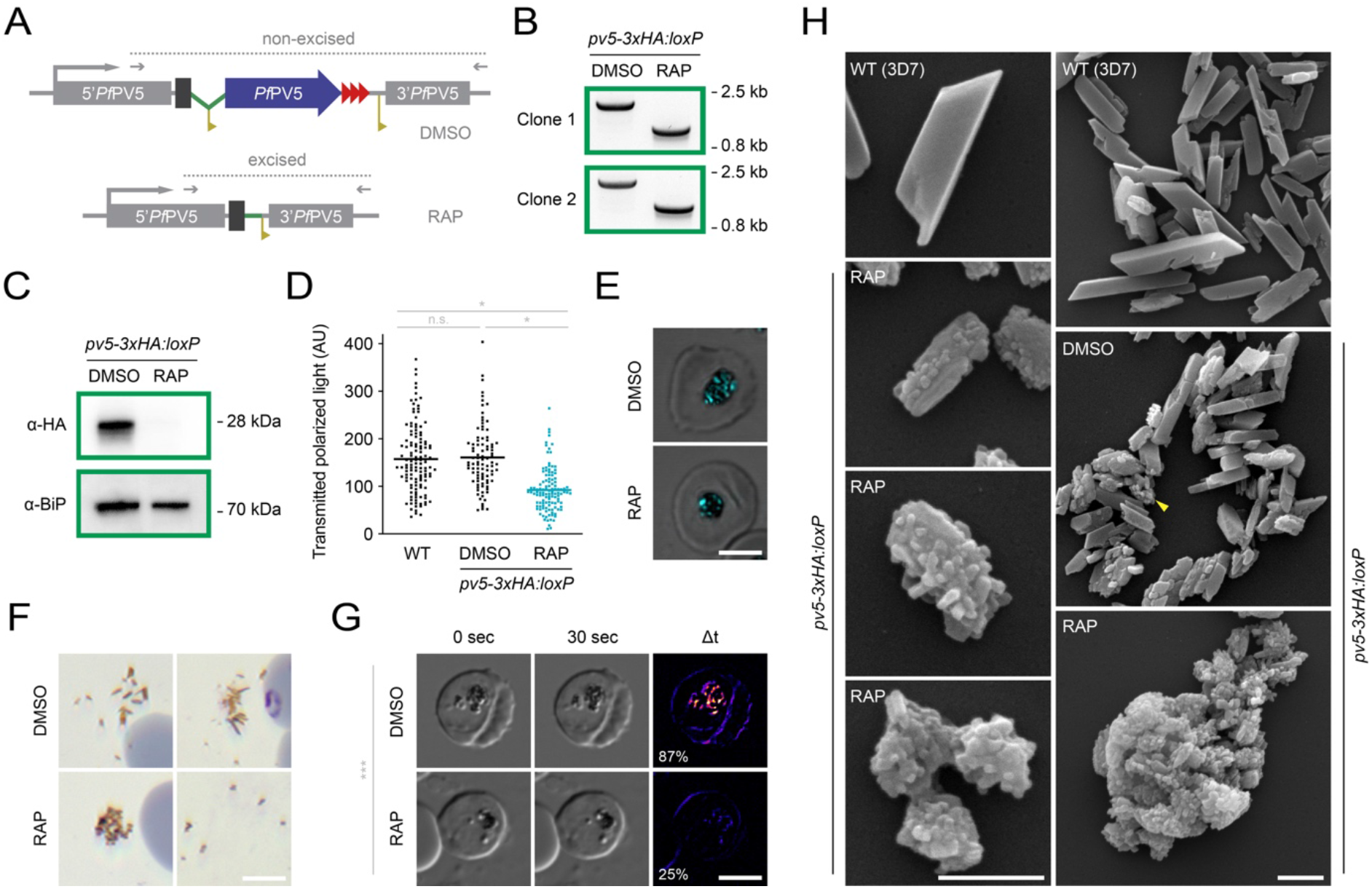
Absence of PV5 causes haemozoin branching in *Plasmodium falciparum*. *(A)* Schematic representation of DiCre-mediated Pf*PV5* excision. The endogenous locus was modified to introduce *loxP* sites (yellow) flanking the majority of the coding sequence of the 3xHA (red)-tagged *Pf*PV5 (blue). The artificial intron is indicated in green. See also Fig. S3*A*. Treatment with rapamycin (RAP) induces Cre recombinase-mediated excision of the *loxP*-flanked sequence which results in truncation of Pf*PV5* leaving behind only the sequence encoding the protein’s signal peptide. Excision-sensitive primer combinations are indicated by arrows and expected diagnostic PCR fragments by dotted lines. (*B*) Diagnostic PCR of the modified Pf*PV5* locus following treatment with dimethyl sulfoxide (DMSO) or RAP, respectively, using the primer combinations depicted in *A*. Results are shown from two independent *pv5-3xHA:loxP* clones. (*C*) Loss of *Pf*PV5 protein. Western blot analysis of parasite extracts following treatment with DMSO or RAP, respectively, using anti-HA and anti-*Pf*BiP primary antibodies. (*D*, *E*) *Pf*PV5 is required for efficient Hz formation. WT and *pv5-3xHA:loxP* parasites were treated with DMSO or RAP, respectively, and visualized by polarization microscopy 36 hours after invasion. (*D*) Quantification of the polarized light intensity. Depicted are values from individual parasites as well as the mean intensity values (bars). n.s., non-significant; *, P<0.05; One-way ANOVA and Tukey’s multiple comparison test. N≥93 parasites from 3 independent experiments. (*E*) Exemplary images of DMSO- and RAP-treated *pv5-3xHA:loxP* parasites. Shown is a merge of polarized light (cyan) and DIC. Bar, 5 μm. (*F*) Abnormal Hz morphology in the absence of *Pf*PV5. Hz released from residual bodies during parasite egress was imaged in Giemsa-stained thin culture smears of DMSO and RAP-treated *pv5-3xHA:loxP* parasites. Note the spreading of elongated Hz crystals in DMSO-treated cultures and the clumping of granular Hz upon RAP treatment (left). Only in very few instances did the crystals detach from one another in RAP-treated cultures (right). Bar, 5 μm. (*G*) Reduced Hz movement in the FV upon loss of *Pf*PV5. Shown are DIC images of live DMSO and RAP-treated *pv5-3xHA:loxP* parasites 36 hours after invasion. Parasites were imaged twice at an interval of 30 seconds (left and centre) and the difference of both images was visualized with pixel-by-pixel intensity subtraction (right). Note the absence of Hz movement in the RAP-treated parasite (see also Movie S1). The percentage of parasites with moving Hz is indicated. ***, P<0.001; Student’s *t*-test. N=4 independent experiments with >300 parasites each. Bar, 5 μm. (*H*)Scanning electron micrographs of Hz purified from WT (3D7) and from DMSO or RAP-treated *pv5-3xHA:loxP* schizonts. Bars, 500 nm.

Treatment of *pv5-3xHA:loxP* parasites with rapamycin led to efficient gene excision and complete loss of *Pf*PV5 protein expression during the same intraerythrocytic cycle (Fig. 4 *B* and *C*, Fig. S3*D*). This did not detectably affect parasite maturation but did result in a modest merozoite invasion defect upon rupture of the Pf*PV5*-null schizonts, reducing parasite replication (Fig. S4). Extended monitoring of the rapamycin-treated parasites indicated an estimated fitness cost of ~40% (Fig. S4*B*). This is in good agreement with a proposed mutagenesis index score of 0.22 from a genome-wide piggy-Bac insertion mutagenesis screen (36). Accordingly, Pf*PV5*, although not essential under standard *P. falciparum* culture conditions, is required for optimal parasite propagation *in vitro*.

To examine the effects of Pf*PV5* ablation on haem biomineralization, Hz was visualized and quantified by polarization microscopy. Rapamycin-treated *pv5-3xHA:loxP* parasites formed only 57% of the Hz observed in WT and DMSO-treated controls and individual crystals appeared to be globular rather than elongated (Fig. 4 *D* and *E*). In the absence of *Pf*PV5, Hz released at parasite egress no longer formed clusters of separate slender crystals but rather appeared as aggregates which only occasionally fell apart into individual globular units (Fig. 4*F*). In good agreement, microscopic inspection of live parasites revealed that the characteristic twirling motion of Hz within the central FV was largely lost upon Pf*PV5* knockout (Fig. 4*G* and Movie S1).

The dramatic abnormalities in *Hz* crystal morphology resulting from ablation of Pf*PV5* were even more evident by SEM analysis. While WT parasites formed crystals of the expected brick-like morphology, individual Hz units from rapamycin-treated *pv5-3xHA:loxP* parasites appeared smaller and more globular (Fig. 4*H* and Table S1). The surfaces of these Hz units were covered in scales and stubby crystal buds. Individual crystals of comparable bud-like dimensions were not observed, indicating a branching rather than an aggregation phenomenon. In some instances, a lower number of crystal buds allowed the visualization of an ordered Hz core, suggesting that branching is initiated from a regular parent crystal (Fig. 4*H*). We detected several morphological intermediates between slightly scaled Hz, highly branched crystal units and fused congregations (Fig. 4*H*). We frequently observed enormous aggregates of spherical proportions, mirroring the shape of the central FV (fig. 4*H*). This suggested that hyperactive crystal branching in the absence of *Pf*PV5 caused individual studded Hz units to stick together and subsequently merge during Hz growth, which might explain the absence of motion in the parasite FV (Fig. 4*H*). Non-rapamycin-treated *pv5-3xHA:loxP* control parasites mainly formed regular Hz crystals, however 27.5% of the crystals exhibited a modest degree of branching (Fig. 4*H* and Table S1). Furthermore, crystal size and aspect ratio were reduced in comparison to WT parasites, suggesting a moderate functional impairment of 3xHA-tagged *Pf*PV5 (Fig. 4*H* and Table S1). Our data demonstrate that PV5 is critical for the efficient sequestration of haem and for the ordered expansion of Hz crystals in *P. falciparum*.

### Efficient haemoglobin processing in the absence of *Pf*PV5

To exclude an indirect effect mediated by defective haemoglobin catabolism, we examined the haemoglobin content of saponin-released parasites. *Pf*PV5-deficient ring stages and trophozoites contained normal quantities of internalized haemoglobin (Fig. 5*A*). Only mature segmented *pv5-3xHA:loxP* schizonts exhibited slightly elevated concentrations of residual haemoglobin upon induction (Fig. 5*B*). However, this was also observed in WT schizonts upon rapamycin treatment, suggesting a minor compound-specific effect. Furthermore, saponin treatment released normal amounts of haemoglobin from schizont-infected erythrocytes in the absence of *Pf*PV5, indicating unaltered haemoglobin ingestion (Fig. 5*B*). As a positive control for perturbation of haemoglobin catabolism, we cultured WT parasites in the presence of the protease inhibitor E64. Vacuolar bloating, as readily detected by E64 treatment, was not visible in rapamycin-treated *pv5-3xHA:loxP* parasites (Fig. 5*C*). We also note that, unlike Pf*PV5* deletion, E64 treatment produced no changes in the architecture of Hz (Fig. 5*D*). In addition, *Pf*PV5-deficient parasites retained normal E64 sensitivity (Fig. 5 *E* and *F*). Combined, these findings suggest that inhibition of haemoglobin catabolism does not directly translate into altered Hz morphology and that *Pf*PV5 is not involved in the overall consumption of host cell cytoplasm.

**Fig. 5.**
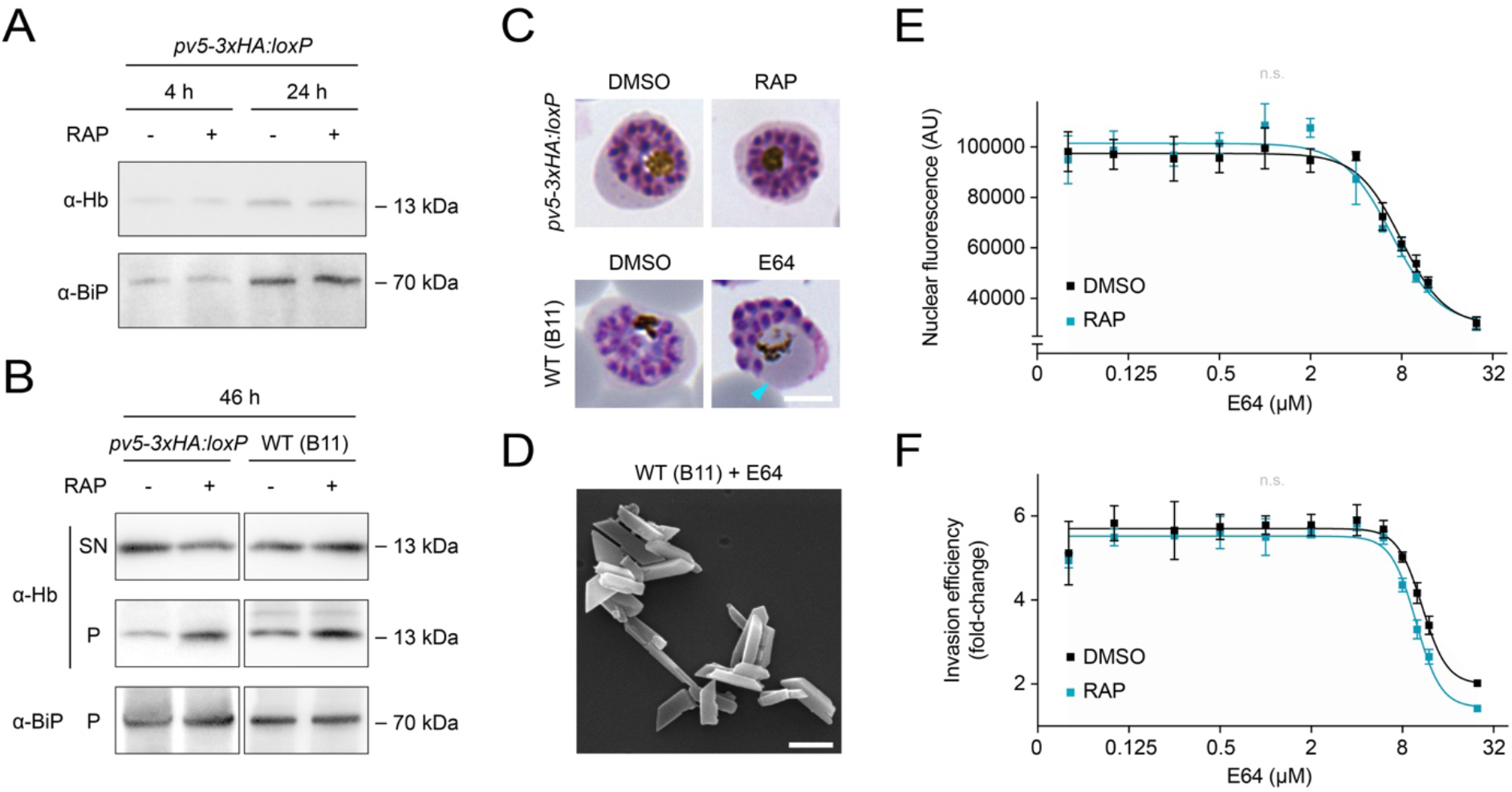
*Pf*PV5 regulates haem sequestration independently from haemoglobin processing. (*A, B*) Normal uptake and digestion of host cell haemoglobin by *Pf*PV5-deficient parasites. (*A*) Western blot analysis of induced and non-induced *pv5-3xHA:loxP* parasites released from their host cells by saponin lysis 4 and 24 hours after invasion. (*B*) 46 hours after invasion, induced and non-induced *pv5-3xHA:loxP* or WT (B11) schizonts were released by saponin treatment. Resultant supernatants (SN) and schizont pellets were isolated. Blots were probed with antibodies directed against human haemoglobin α (Hb) and *Pf*BiP. Note an increase in intraparasitic haemoglobin upon rapamycin treatment (RAP) in both *pv5-3xHA:loxP* and WT (B11) schizonts. (*C*) *Pf*PV5-deficient schizonts exhibit no vacuolar bloating. Shown are *pv5-3xHA:loxP* parasites treated from the ring stage onward with dimethyl sulfoxide (DMSO) or RAP (top) and *Plasmodium falciparum* WT parasites treated from 24 hours post invasion onward with DMSO or 21.7 μM E64 (bottom). Cyan arrowhead, bloated food vacuole. Bar, 5 μm. (*D*) Inhibition of haemoglobin catabolism does not result in abnormal Hz morphology. Shown is a scanning electron micrograph of Hz crystals isolated from the E64-treated *P. falciparum* WT parasites shown in *C*. Bar, 500 nm. (*E, F*) *Pf*PV5-deficient parasites display unaltered sensitivity towards E64. DMSO and RAP-treated *pv5-3xHA:loxP* parasites were grown in various concentrations of E64 from the ring stage onward. (*E*) Nuclear SYBR Green fluorescence 44 hours after invasion. (*F*) Invasion efficiency under static conditions following a 36-hour incubation in the presence of E64 and subsequent inhibitor washout. Mean values +/-SD are shown. n.s., non-significant; fitting of IC_50_ values following non-linear regression, N=3.

### Pb*PV5* expression influences antimalarial drug sensitivity *in vivo*

In the light of our evidence implicating PV5 in Hz formation, we next tested whether parasites with affected PV5 function display altered sensitivity towards chloroquine, a 4-aminoquinoline which is thought to inhibit haem biomineralization in *Plasmodium* (22–25). The absence of *Pf*PV5 did not detectably alter chloroquine sensitivity in cultured *P. falciparum* parasites (Fig. 6*A*). In stark contrast to this, however, the *P. berghei* promoter swap mutants responded to drug treatment slightly earlier and disappeared from the circulation much more rapidly than WT parasites (Fig. 6*B*). A similar phenotype was observed upon treatment of infected mice with the artemisinin derivative artesunate, also previously been implicated in haem sequestration, although this remains contentious (37–41) (Fig. 6*C*). Most surprisingly, we also observed marked hypersensitivity of the *P. berghei* mutants towards atovaquone, a compound that targets the parasite’s mitochondrial electron transport chain (Fig. 6*D*), as well as a slight but non-significant increase in sensitivity towards the antifolate sulfadoxin (Fig. 6*E*). The relative survival levels of *pv5::5’ptex88* and *pv5::5’hsp101* on the fourth day of drug treatment suggested the greatest degree of hypersensitivity towards chloroquine (2.1% and 1.4% of WT survival, respectively), followed by atovaquone (3.5% and 8.6%) and artesunate (4.4% and 9.9%), and eventually sulfadoxin (20.5% and 27.8%). An unrelated slow-growing *P. berghei* mutant deficient in the maintenance of the mitochondrial membrane potential displayed normal sensitivity towards atovaquone using the same internally controlled assay (42), indicating that reduced parasite multiplication is unlikely to cause drug hypersensitivity. Collectively, these results suggest that interference with *PV5* expression critically enhances vulnerability of the parasite towards drug-mediated insult during *in vivo* blood infection.

**Fig. 6.**
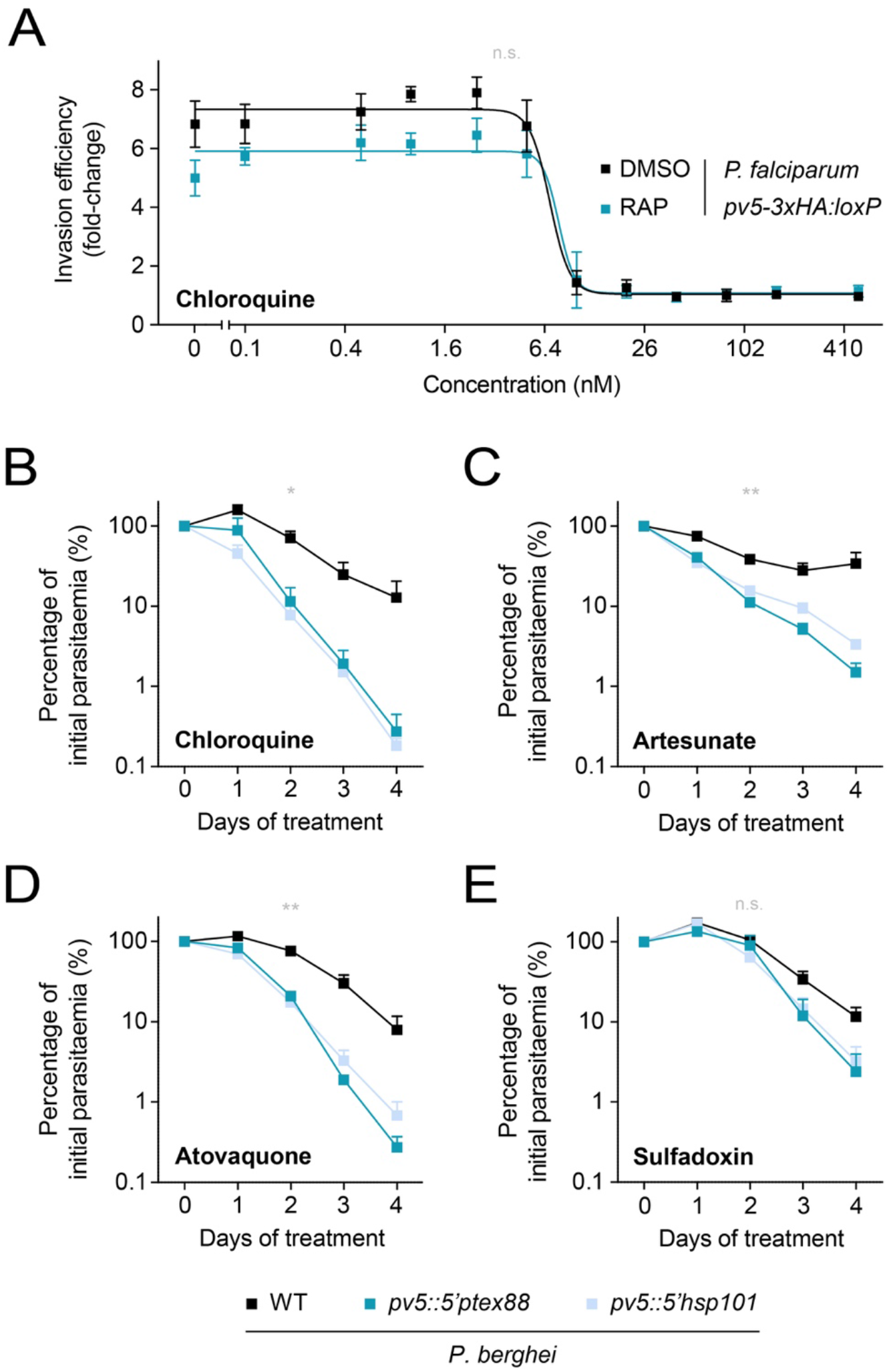
Targeting PV5 function results in parasite hypersensitivity towards antimalarial drugs *in vivo*. (*A*) PV5-deficient *P. falciparum* parasites retain normal sensitivity towards chloroquine. Shown is a dose-response analysis of DMSO- and RAP-treated *pv5-3xHA:loxP* parasites. Static ring stage cultures were treated with varying chloroquine concentrations and the transition into the following intraerythrocytic cycle was quantified. Depicted are mean values +/-SD. n.s., non-significant; fitting of IC50 values following non-linear regression, N=3. (*B*-*E*) Enhanced drug susceptibility of the Pb*PV5* mutants *in vivo*. 5 x 10^6^ mCherry-fluorescent *P. berghei* WT parasites were injected into SWISS mice together with 5 x 10^6^ GFP-fluorescent *pv5::5’ptex88* or *pv5::5’hsp101* parasites, respectively. From day three onward, mice were treated with curative doses of (*A*) chloroquine (288 mg/l in drinking water, *ad libitum)*, (*B*) artesunate (50 mg/kg body weight, i.p.), (*C*) atovaquone (1.44 mg/kg body weight, i.p.) or (*D*) sulfadoxin (1.4 g/l in drinking water, *ad libitum)* and the respective parasitaemias were determined daily by flow cytometry of peripheral blood (50). Values are normalized to the parasitaemias on day 0 of treatment. Shown are mean values +/-SEM. n.s., non-significant; *, P<0.05; **, P<0.01; Two-way ANOVA. N=3 independent infections.

## DISCUSSION

In this work, we have demonstrated that Hz formation in malaria parasites involves a secreted calycin family member called PV5. While transcriptional deregulation of *PV5* in *P. berghei* resulted in protracted Hz elongation along a pre-existing crystal plane, multidirectional branching was observed in the complete absence of PV5 in *P. falciparum*. Species differences aside, the disparity between those two phenotypes can be explained by the unique transcriptional dynamics observed in the *P. berghei* mutants. Here, the hyperactive crystal elongation, that follows an initial phase of normal Hz growth, coincides with a substantial increase in Pb*PV5* transcript abundance during late parasite development. In their physiological context, PTEX88 and HSP101 are co-expressed as members of the same protein complex with HSP101 being more abundant than PTEX88 (34, 43), as supported by our qPCR analysis. Thus, the increased phenotypic severity in the *pv5::5’hsp101* mutant over the *pv5::5’ptex88* mutant indicates that the deficiencies in Hz formation can be attributed to late overexpression of PV5. Together with the chaotic Hz crystal branching observed in PV5-deficient *P. falciparum* parasites, this leads us to propose that PV5 acts as a facilitator of unidirectional Hz extension.

It is interesting to speculate on the mechanisms by which PV5 may partake in haem biomineralization. In laboratory crystallization experiments, the extent of crystallographic mismatch branching is highly dependent on the degree of solute supersaturation, with high levels favouring the generation of novel crystal nuclei on the surface of the parent crystal (44). By contrast, lower degrees of supersaturation usually promote the expansion of pre-existing crystals (44). It is conceivable that PV5 might reduce the extent of haem supersaturation by binding haem or haematin dimers, thereby moderating *de novo* Hz nucleation and promoting unidirectional crystal elongation. This could explain the Hz branching upon loss of PV5 in *P. falciparum* as well as the prolonged crystal elongation and reduced Hz production upon late upregulation of *PV5* in the *P. berghei* mutants. In support of this model, some lipocalin family members are known to specifically bind the haem degradation product biliverdin (45–47). Such interactions are unlikely to occur within the predicted PV5 β-barrel due to spatial constraints. However, although experimental validation for this is currently lacking, potential ligand binding sites might be located at the predicted prominently exposed loop between the barrel’s β-strands 5 and 6 (Fig. 1*B*) or at the extended amino-terminus.

The crystal branching characterising PV5-deficient *P. falciparum* parasites could also be elicited by non-haem impurities adsorbing onto the crystal surface where they would generate novel nucleation sites (44). In this alternative functional model, PV5 could act to bind these impurities to create a vacuolar environment permissive for proper biomineralization. *Pf*PV5-deficient parasites maintain an intact FV with a trans-vacuolar proton gradient as indicated by staining with Lysosensor Blue DND-167 (Fig. S5*A*). In the absence of indications for FV membrane damage, differences in leakage of impurities from the parasite cytoplasm can be dismissed. However, we cannot rule out the possibility that specific ions or organic compounds are more abundant in the vacuolar matrix of PV5-deficient parasites.

While the exact biophysical mechanisms that govern PV5-mediated Hz morphogenesis remain to be delineated, our results support a model of conventional crystallization within the aqueous milieu of the parasite’s digestive compartments. The distinct changes in crystal morphology and branching behaviour in the *PV5* mutants concur with established mineralogical phenomena and are difficult to reconcile with a lipid or protein-mediated polymerization scenario.

The striking dual localization pattern of PV5 is suggestive of initial secretion into the PV followed by endocytic uptake. Since we ruled out an involvement of PV5 in haemoglobin ingestion and catabolism, the transient vacuolar swelling in the *P. berghei* mutants likely reflects a secondary effect mediated by the grave deficiencies in haem sequestration.

Our previous experiments had indicated that Pb*PV5* is essential for the asexual blood stage development of *P. berghei* (27), and the deregulation of Pb*PV5* transcription in the mutants described here indeed led to a striking fitness cost during *in vivo* infection. By contrast, complete ablation of Pf*PV5* in *P. falciparum* only resulted in a moderate fitness loss *in vitro*. The apparent dispensability of Pf*PV5* is therefore puzzling, but can be resolved by the notion that *in vitro* culture does not necessarily reflect all adversities encountered during host infection. For instance, Hz-mediated stiffening of the residual body in the absence of PV5 might hinder the passage of infected cells through capillaries or through the inter-endothelial slits of the spleen, a scenario that would result in parasite elimination only during *in vivo* infection. Similarly, we observed enhanced drug susceptibility only in the *P. berghei* mutants. Thus, it appears plausible, that any imbalance in haem biomineralization in the absence of PV5 is compensated for under optimal culture conditions and unfolds its adverse effects only during *in vivo* infection. We thus propose that the molecular mechanism by which PV5 regulates Hz formation might involve host factors that are not encountered in a cell culture setting.

The observation of enhanced drug susceptibility in the Pb*PV5* mutants is in agreement with the notion that aberrant haem biomineralization chemo-sensitizes parasites to partner drugs. Although oxidative stress and lipid peroxidation remained unchanged upon deletion of Pf*PV5* (Fig. S5*B*), the reduced Hz formation efficiency in the *PV5* mutants suggests elevated haem concentrations within the parasite mediating the fitness loss and drug hypersensitivity.

Together, we provide the first conclusive evidence for a parasite factor mediating Hz formation *in vivo*, called PV5. Since this *Plasmodium-encoded* member of the calycin superfamily also governs the parasites’ vitality and susceptibility towards drug-mediated insult during blood infection, our observations reinforce Hz formation as an excellent pathway for therapeutic intervention. The investigation of the malaria parasite’s haem detoxification machinery *in vivo*, as exemplified herein for PV5, will significantly improve our understanding of this unique haem biomineralization process and holds great promise for the development of novel malaria intervention strategies.

## MATERIALS & METHODS

### Structure homology modelling

Structure homology modelling was performed using the SWISS-MODEL server (48). *Pf*PV5 (residues 35-214) was aligned to the experimentally validated structure of *Ec*Blc (residues 27-175, PDB ID: 3MBT) (49), resulting in a GMQE value of 0.39 and a QMEAN value of −5.48. Modelling was confirmed with I-TASSER (50), which also identified *Ec*Blc (PDB ID: 2ACO) (51) as the most closely related structural analogue of *Pf*PV5 with a TM score of 0.75. Due to a lack of sequence similarity, the structure of the extended *Pf*PV5 amino-terminus was not modelled.

### *P. berghei* cultivation

*P. berghei* parasites were propagated in SWISS mice under constant drug pressure with pyrimethamine (70 mg/l in drinking water, ingested *ad libitum*, MP Biomedicals) to avoid homology-induced reversion of the promoter swap mutants to the original WT genotype. This was routinely checked by diagnostic PCR of genomic DNA as shown in Figures S2 *A* and *B*. Drug pressure was withdrawn 5 days prior to experimentation to avoid secondary effects of pyrimethamine treatment. Pyrimethamine-resistant Berred WT parasites (52) were treated accordingly. All infection experiments were carried out in strict accordance with the German ‘Tierschutzgesetz in der Fassung vom 22. Juli 2009’ and the Directive 2010/63/EU of the European Parliament and Council ‘On the protection of animals used for scientific purposes’. The protocol was approved by the ethics committee of the Berlin state authority (‘Landesamt fur Gesundheit und Soziales Berlin’, permit number G0294/15).

*P. berghei* growth was determined with the previously described intravital competition assay (52). In short, 500 mCherry-fluorescent Berred WT and 500 GFP-fluorescent mutant blood stage parasites were co-injected intravenously and parasitaemia was analysed daily by flow cytometry. For drug sensitivity assays, 5×10^6^ WT and 5×10^6^ mutant parasites were co-injected intravenously. Drug treatment as well as daily flow cytometric parasite detection were commenced three days later (43). Mice were treated with curative doses of chloroquine (288 mg/l in drinking water, ingested *ad libitum*, Sigma Aldrich), atovaquone (1.44 mg/kg body weight per day injected intraperitoneally; GlaxoSmithKline), artesunate (50 mg/kg body weight per day injected intraperitoneally; Sigma Aldrich) or sulfadoxin (1.4 g/l in drinking water, ingested *ad libitum*; Sigma Aldrich). Mixed blood stages and schizonts were purified by nycodenz gradient centrifugation (53).

### *P. falciparum* cultivation

*P. falciparum* parasites were propagated in type AB+ human red blood cells at 90% N_2_, 5% CO_2_ and 5% O_2_ at 37°C in RPMI 1640 containing AlbuMAXII (Thermo Fisher Scientific) supplemented with 2 mM L-glutamine. Parasites were routinely synchronized using a combination of percoll gradient centrifugation and sorbitol lysis and were treated with 100 nM rapamycin, various concentrations of E64 or equivalent volumes of dimethyl sulfoxide (DMSO) from the early ring stage onward. Growth assays were performed as described previously (54) and parasitaemia as well as DNA content were measured by flow cytometry using the nuclear dye SYBR Green (1:10,000; Thermo Fisher Scientific). For invasion assays, mature schizonts were incubated at 2% initial parasitaemia under static or shaking (120 rpm) conditions, as confirmed by flow cytometry. Parasitaemia was measured again 24 hours after inoculation and the fold-change was calculated. For drug- and inhibitor-response analyses, *pv5-3xHA:loxP* ring stage cultures at 0.5 – 2% parasitaemia were treated with varying concentrations of chloroquine or E64 in the presence of DMSO or rapamycin in a 96 well format. Parasitaemia was determined three days later and the fold-change was calculated. E64 was washed out after 36 hours to allow for reinvasion. In addition, nuclear SYBR Green fluorescence was determined by flow cytometry following 44 hours of continuous E64 treatment.

### Generation and validation of transgenic *Plasmodium* parasites

To generate the Pb*PV5* promoter swap mutants, the amino-terminal sequence of Pb*PV5* (PBANKA_0826700) was PCR-amplified and cloned into the B3D+ vector using BamHI and SacII. The promoters of *PbPTEX88* (PBANKA_0941300) or Pb*Hsp101* (PBANKA_0931200) were then amplified and cloned in front of the start codon using BamHI (Figure S1*A*). Vectors were linearized with BstBI and transfected into GFP-fluorescent *P. berghei* Bergreen WT parasites, using standard protocols (53, 55, 56). Transgenic parasites were selected for with pyrimethamine and isolated by limiting dilution cloning. Pb*PV5* transcript abundance in the WT and the promoter swap mutants was measured by quantitative real-time PCR (qPCR) and normalized to *Pb*18S rRNA.

The conditional Pf*PV5* knockout line was generated using established Cas9-mediated techniques. In short, *P. falciparum* 3D7 parasites constitutively expressing DiCre (B11 line) were co-transfected with a pDC2 guide plasmid inducing Cas9-mediated double strand cleavage of the Pf*PV5* locus (PF3D7_0925900), together with a linearized repair template (57–59) (Figure S3*A*). The template was generated by gene synthesis and contained 5’ and 3’ homology arms and a re-codonised 3xHA-tagged Pf*PV5* sequence. The endogenous intron of Pf*PV5* was replaced with an artificial intron containing a *loxP* site (58). A second *loxP* sequence was introduced behind the Pf*PV5* stop codon. For live imaging of *Pf*PV5, mCherry was cloned into the conditional knockout vector in frame with the re-codonised Pf*PV5* sequence using AatII. Transgenic parasites were selected for with WR99210 and cloned by limiting dilution, making use of a previously developed plaque assay (60). Primers used for molecular cloning, diagnostic PCR, and qPCR are indicated in Figures 4*A*, S1*A* and S3*A* as well as in Table S2.

### Fluorescence microscopy

The transgenic *P. berghei* parasite line *pv5-tag-GFP*^PV^ (27) was imaged live using an AxioImager Z2 epifluorescence microscope equipped with an AxioCam MR3 camera (Zeiss). For mechanical parasite expansion, 1-2 μl of infected blood were incubated under a 22 x 40 mm coverslip for several minutes until erythrocyte lysis became apparent. *P. falciparum* parasites were imaged on an Eclipse Ni light microscope (Nikon) fitted with a C11440 digital camera (Hamamatsu). Immunofluorescence analysis was performed with *P. falciparum pv5-3xHA:loxP* parasites that were fixed in 4% formaldehyde using rat anti-HA (1:500; Sigma Aldrich) and rabbit anti-SERA5 (1:500) (54) primary antibodies in combination with appropriate fluorophore-coupled secondary antibodies (1:1,000; Thermo Fisher Scientific). Stainings with Lysosensor Blue DND-167 (Thermo Fisher Scientific), BODIPY 581/591 C11 (Image-iT Lipid Peroxidation Kit, Thermo Fisher Scientific) and CellROX Green (Thermo Fisher Scientific) were performed according to the manufacturer’s instructions.

### Scanning electron microscopy

Hz was isolated from nycodenz (*P. berghei*) or percoll (*P. falciparum*) gradient-purified infected red blood cells. Cells were lysed in water at room temperature for 20 minutes, followed by a 10-minute centrifugation step at 17,000 x *g*. The pellet was resuspended in 2% SDS in water and centrifuged as above. Three more washing steps with 2% SDS were then followed by three washing steps with distilled water, before the crystals were resuspended and transferred onto round glass cover slips (12 mm), where they were dried. Cover slips were mounted on SEM specimen stubs, sputter-coated, and then imaged on a LEO 1430 (Zeiss) or on a Quanta FEG 250 scanning electron microscope (Thermo Fisher Scientific).

### Transmission electron microscopy

Infected erythrocytes were initially fixed in 2.5% glutaraldehyde, embedded in beads of 2% agarose, fixed and contrasted with 1% osmium tetroxide, and further contrasted *en bloc* using 0.5% uranyl acetate. Following dehydration in a graded series of ethanol and propylene oxide, beads were embedded in epoxy resin and cured at 60°C for at least 24 hours. 60 nm sections were made with a Reichert Ultracut S ultramicrotome (Leica) using a diamond knife. Sections were retrieved on copper hexagonal mesh grids, and stained with 2% uranyl acetate and Reynold’s lead citrate before imaging on an EM 900 transmission electron microscope (Zeiss) equipped with a wide-angle slow-scan 2K CCD camera (Tröndle Restlichtverstärkersysteme).

### Electron diffraction

Purified Hz was added to glow-discharged Lacey carbon films on 400 mesh copper grids which were then transferred to a Vitrobot Mark IV plunge freezer (Thermo Fisher Scientific) with 100% humidity at 7°C. The grids were blotted for 3 seconds with blot force 1 and plunge frozen in liquid ethane cooled by liquid nitrogen. Electron diffraction data were collected on a Talos cryo-electron microscope (Thermo Fisher Scientific) operated at 200 keV equipped with a hybrid pixel Timepix detector (512 x 512 pixels, 55 x 55 μm pixel size; Amsterdam Scientific Instruments). Still and rotation (70°) datasets were collected with a beam size of 2 μm. The recording time varied between 15 and 100 seconds. To determine the resolution, a powder pattern of an aluminum diffraction standard was recorded. Since the Hz crystals had a strong tendency to stick together, we measured diffraction data of crystal clusters. For each individual data set, we determined the location of the central electron beam and shifted the patterns to make the beams coincide. Since crystal clustering prevented indexing of the diffraction data, the radial median and maximum intensities were determined as a function of resolution. Hereafter, the WT and *pv5::5’hsp101* datasets were averaged and normalized to the local background median intensity at 0.18 Å^-1^.

### Subcellular fractionation and immunoblotting

*P. falciparum* parasites were released from erythrocytes by treatment with 0.15% saponin in PBS. Murine erythrocytes infected with the *pv5-tag-GEP*^PV^ or *exp2-mCherry P. berghei* lines (64) were purified on a nycodenz gradient and lysed hypotonically for 1 hour on ice in 10 mM TRIS-HCl, pH 7.5. *P. berghei* lysates were spun 50 minutes at 100,000 × *g*. Membrane pellets were resuspended in 0.1 M Na2CO3 in PBS or in 1% Triton X-100 in PBS, respectively, and spun 50 minutes at 100,000 ×*g*. Proteins were separated on SDS-polyacrylamide and transferred onto a nitrocellulose membrane. Western blotting was performed using rat anti-mCherry (1:5,000; ChromoTek), chicken anti-GFP (1:5,000; Abcam), rat anti-HA (1:1,000, Sigma Aldrich), rat anti-*Pf*BiP (1:1,000) (62) and rabbit anti-human-haemoglobin-α primary antibodies (1:1,000, Abcam) followed by chemiluminescence detection with horseradish peroxidase-coupled secondary antibodies (1:10,000; Sigma Aldrich, or 1:5,000, Jackson ImmunoResearch).

### Quantitative haemozoin analysis

Hz was visualized and quantified by microscopy of methanol-fixed infected red blood cells. Hz from *P. berghei* was analysed by reflection contrast polarized light microscopy using a Leica DMR widefield microscope equipped with a ProgRes MF camera (Jenoptik) and the POL filter set 513813 (Leica). Hz from *P. falciparum* was analysed by transmitted polarized light (488 nm) microscopy using an LSM 710 confocal microscope (Zeiss) equipped with a crossed polarizer in the condenser. Cellular granularity was approximated by quantification of side scattered light using an LSRFortessa flow cytometer (BD Biosciences). Hz crystal dimensions were analysed using FIJI. Due to the variation in the *P. berghei* mutant’s crystal width, individual crystals were divided into nine evenly spaced segments along the dominant axis. The width of each segment was determined and the values were averaged. For the *pv5::5’hsp101* parent crystal and the crystal outgrowth, transects were drawn through the central axis of either structure and their shared angle was determined. Hz movement within the FV of *pv5-3xHA:loxP* parasites was imaged live 36 hours following treatment with DMSO or rapamycin, respectively.

## AUTHOR CONTRIBUTIONS

JMM conceived the study and performed all experiments requiring genetic manipulation and cultivation of *P. berghei* and *P. falciparum* parasites. JMM, BD, AB, PM, TS and LC contributed to Hz analysis by SEM and TEM. TBB, EvG and JPA performed and analysed electron diffraction experiments. KM and MJB contributed to data interpretation. The manuscript was written by JMM with input from TBB, EvG, JPA, KM and MJB.

## ACKNOWLEDGEMENTS

We thank Volker Brinkmann and Christian Goosmann (Max Planck Institute for Infection Biology, Berlin) as well as Kurt Anderson (The Francis Crick Institute, London) for assistance with polarization microscopy. We also thank Darren Flower (Aston University, Birmingham) and Leslie Leiserowitz (Weizmann Institute of Science, Rehovot) for fruitful discussion. This work was supported by a stipend from the Deutsche Forschungsgemeinschaft (DFG, project number 419345764 to JMM) and in part by funding to JMM, LC, AB and MJB from the Francis Crick Institute, which receives its core funding from Cancer Research UK (FC001043), the UK Medical Research Council (FC001043), and the Wellcome Trust (FC001043). JMM, BD, PM, TS and KM were also supported by the Humboldt University and by the Alliance Berlin Canberra ‘‘Crossing Boundaries: Molecular Interactions in Malaria’’, which is co-funded by a grant from the DFG for the International Research Training Group (IRTG) 2290 and the Australian National University. We further acknowledge funding from the Swiss National Science Foundation (project 31003A_17002 to TBB and project 200021_165669 to TBB and JPA). We thank the Center for Cellular Imaging and NanoAnalytics for support and use of the cryo-electron microscope.

---

**Fig. S1.**
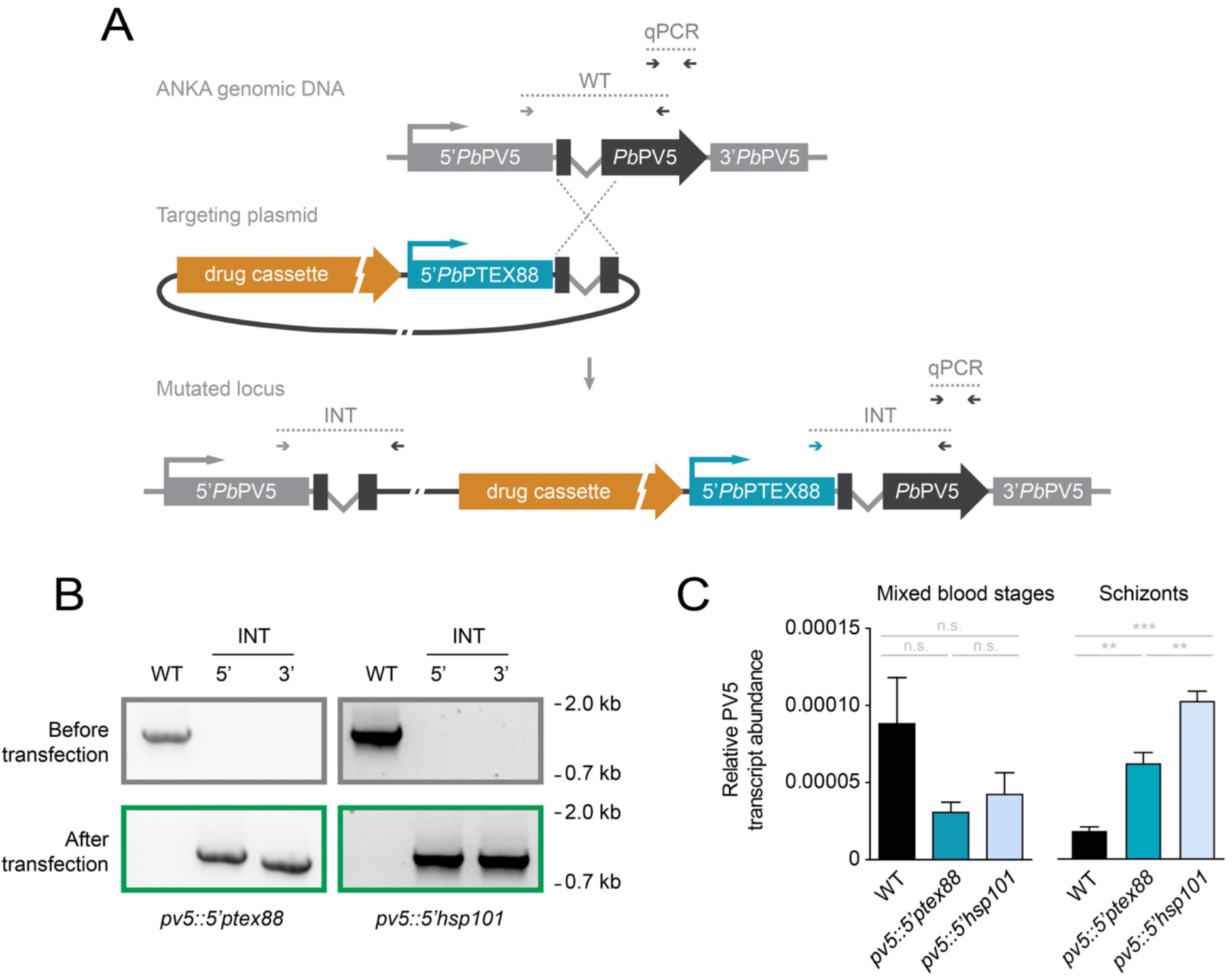
Generation and validation of Pb*PV5* promoter swap mutants. (*A*) Genetic strategy to exchange the endogenous promoter (light grey) of Pb*PV5* (dark grey) using single homologous recombination. The endogenous Pb*PV5* locus was targeted with an insertion plasmid containing the promoter sequence of *PbPTEX88* (shown, blue) or *PbHSP101* (not shown) fused to the amino-terminal sequence of Pb*PV5* as well as the drug-selectable hDHFR-yFcu cassette (orange). Successful insertion yields parasites expressing full length Pb*PV5* from a heterologous promoter and a non-functional amino-terminal fragment from the endogenous promoter, encoding the *Pb*PV5 signal peptide. Primer combinations for wild-type (WT) and integration-specific reactions (INT) as well as for quantitative real-time PCR (qPCR) are indicated by arrows and expected amplicons by dotted lines. (*B*) Diagnostic PCR of the WT locus (top) and of the drug-selected and isolated Pb*PV5* mutants (bottom) using the primer combinations depicted in *A*. (*C*) Dynamic changes in Pb*PV5* transcription upon promoter swap. Mixed blood stages and mature segmented schizonts were purified from mouse blood or from *in vitro* culture, respectively, and subjected to qPCR using primers targeting a carboxy-terminal portion of Pb*PV5*, as depicted in *A*. Pb*PV5* transcript abundance was normalized to *Pb*18S rRNA. Shown are mean values +/-SEM. n.s., non-significant; **, P<0.01; ***, P<0.001; One-way ANOVA and Tukey’s multiple comparison test. N=6 for mixed blood stages and 3 for schizonts.

**Fig. S2.**
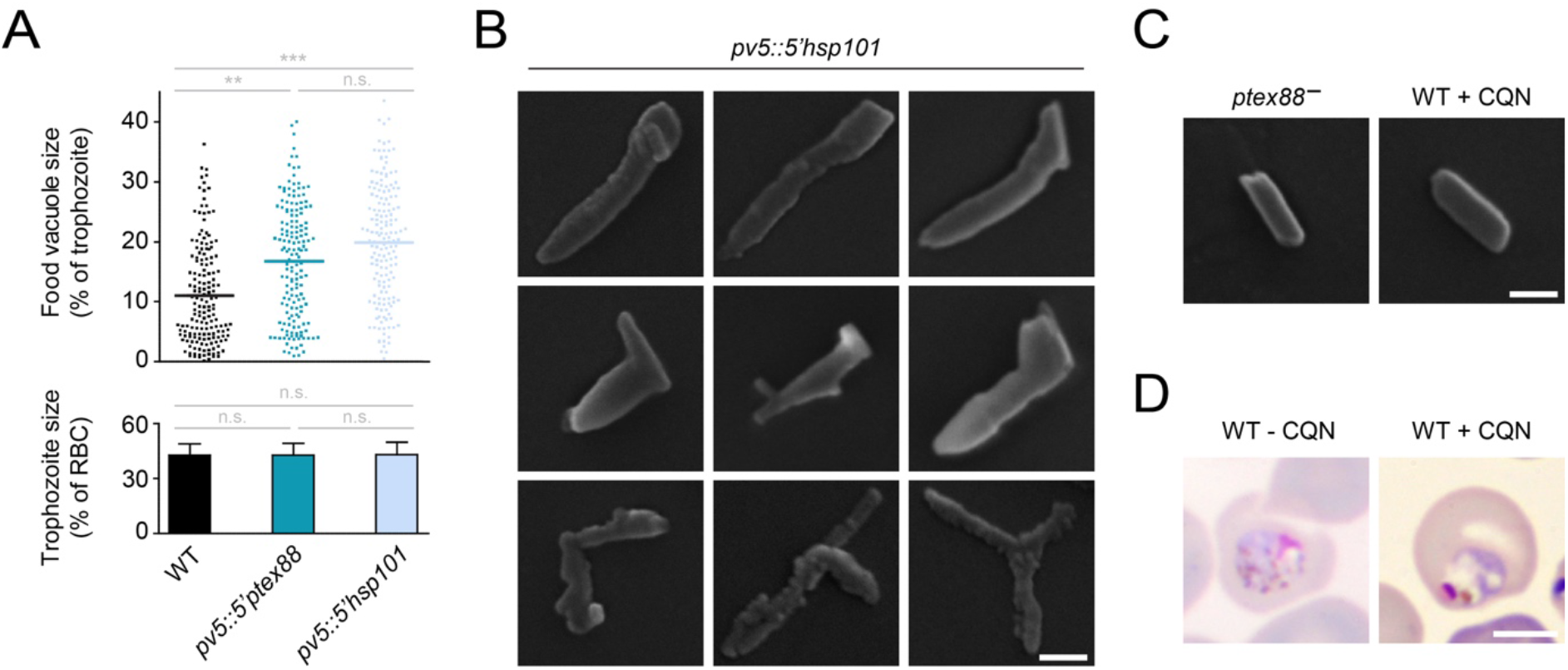
Vacuolar dilation and defective haemozoin formation in the Pb*PV5* mutants. (*A*) Quantification of vacuolar dilation. The translucent area observed in Giemsa-stained trophozoites corresponding to the FV was measured microscopically and expressed as the percentage of the entire trophozoite area (upper graph). Depicted are individual and mean values (bars). Only trophozoites of identical size were analysed (lower graph). Shown are mean values +/-SD. n.s., non-significant; **, P<0.01; ***, P<0.001; One-way ANOVA and Tukey’s multiple comparison test. N=165 trophozoites from 5 independent infections. (*B*) A selection of Hz crystals generated by *pv5::5’hsp101* parasites as visualized by scanning electron microscopy. Bar, 100 nm. (*C*, *D*) Hz morphology is not affected by slow parasite growth or mortality. (*C*) Crystals were extracted from slow-growing *PTEX88* knockout parasites (left) and from WT parasites treated with curative doses of chloroquine (CQN, 288 mg/l in drinking water, *ad libitum)* (right) and were visualized by scanning electron microscopy. Bar, 100 nm. (*D*) Morphology of untreated (left) and CQN-treated WT parasites (right) as shown by Giemsa staining. Note the pigment clumping and vesiculation in the dying parasite. Bar, 5 μm.

**Fig. S3.**
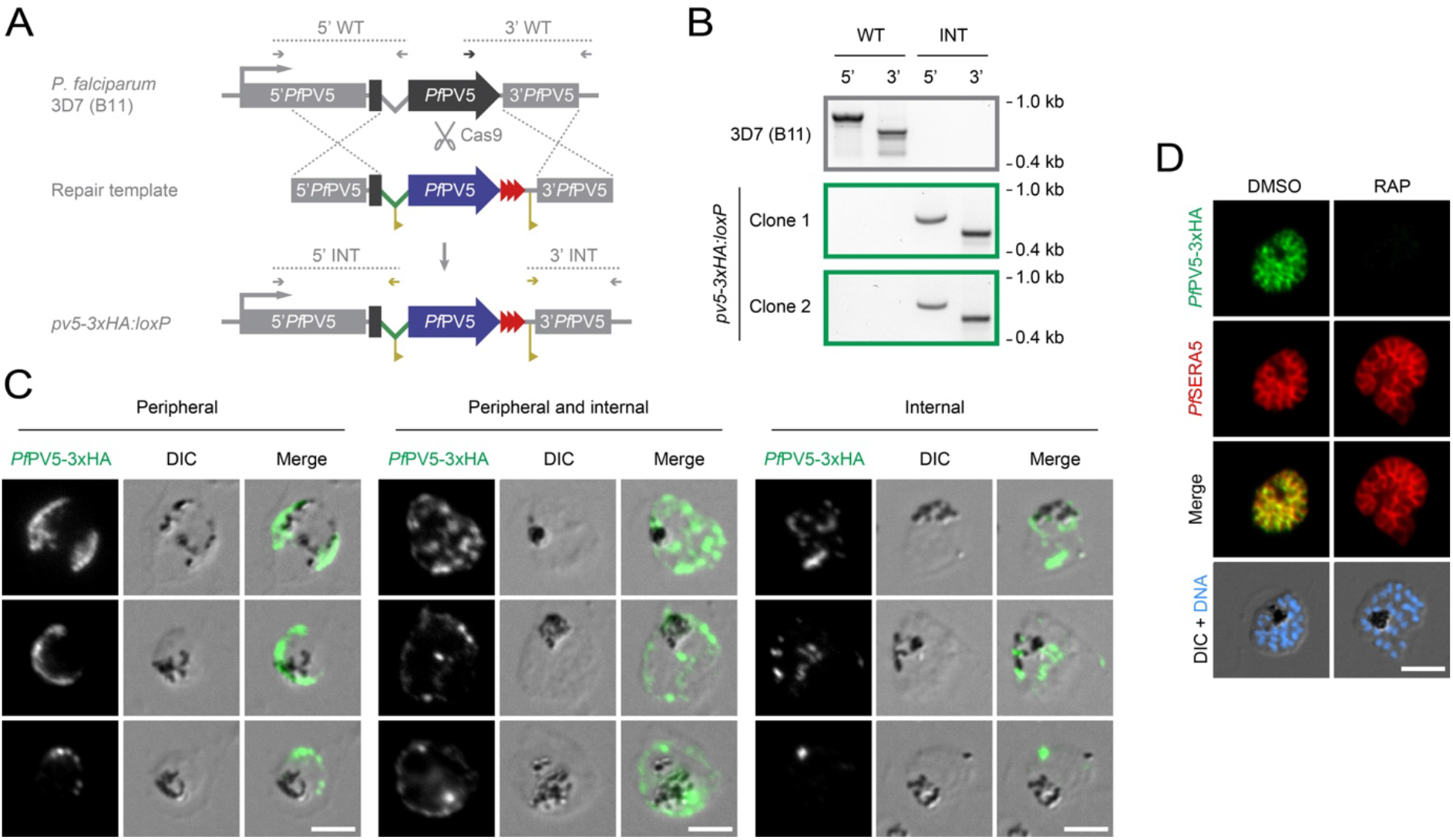
Generation and validation of conditional Pf*PV5* knockout parasites. (*A*) Genetic strategy for the generation of a conditional Pf*PV5* knockout line. The endogenous Pf*PV5* locus (dark grey) was targeted by Cas9-mediated double strand cleavage and repaired by homologous recombination with a synthetic template containing 5’ and 3’ homology arms (light grey), a 3xHA-tagged (red) re-codonised version of Pf*PV5* (blue) and *loxP* sequences (yellow) within the artificial intron (green) and behind the stop codon. Wild-type (WT) and integration-specific primer combinations (INT) are indicated by arrows and expected amplicons by dotted lines. Note that a second conditional knockout line was generated expressing *Pf*PV5 fused to mCherry (Fig. 1*E*). (*B*) Diagnostic PCR of the recipient *P. falciparum* 3D7 (B11) line (top) and of two isolated *pv5-3xHA:loxP* clones (bottom) using the primer combinations depicted in *A*. (*C*) Dual localization of 3xHA-tagged *Pf*PV5. Immunofluorescence analysis was performed using anti-HA primary antibodies. Depicted are exemplary *pv5-3xHA:loxP* trophozoites and young schizonts demonstrating localization of 3xHA-tagged *Pf*PV5 exclusively to the parasitophorous vacuole (PV, left), to the PV and internal parasite structures (centre) or to intraparasitic structures only (right). Shown are the signal of tagged *Pf*PV5 (green, left columns), differential interference contrast images (DIC, centre columns) and a merge (right columns). (*D*) Loss of *Pf*PV5 protein. Immunofluorescence analysis of *pv5-3xHA:loxP* schizonts was performed with primary antibodies directed against HA and the PV protein *Pf*SERA5. Shown are the individual signals of 3xHA-tagged *Pf*PV5 (green, first row) and *Pf*SERA5 (red, second row), a merge of both signals (third row) as well as a merge of DIC with Hoechst 33342 nuclear stain (blue, fourth row) following treatment with DMSO (left) or RAP (right), respectively. Bars, 5 μm.

**Fig. S4.**
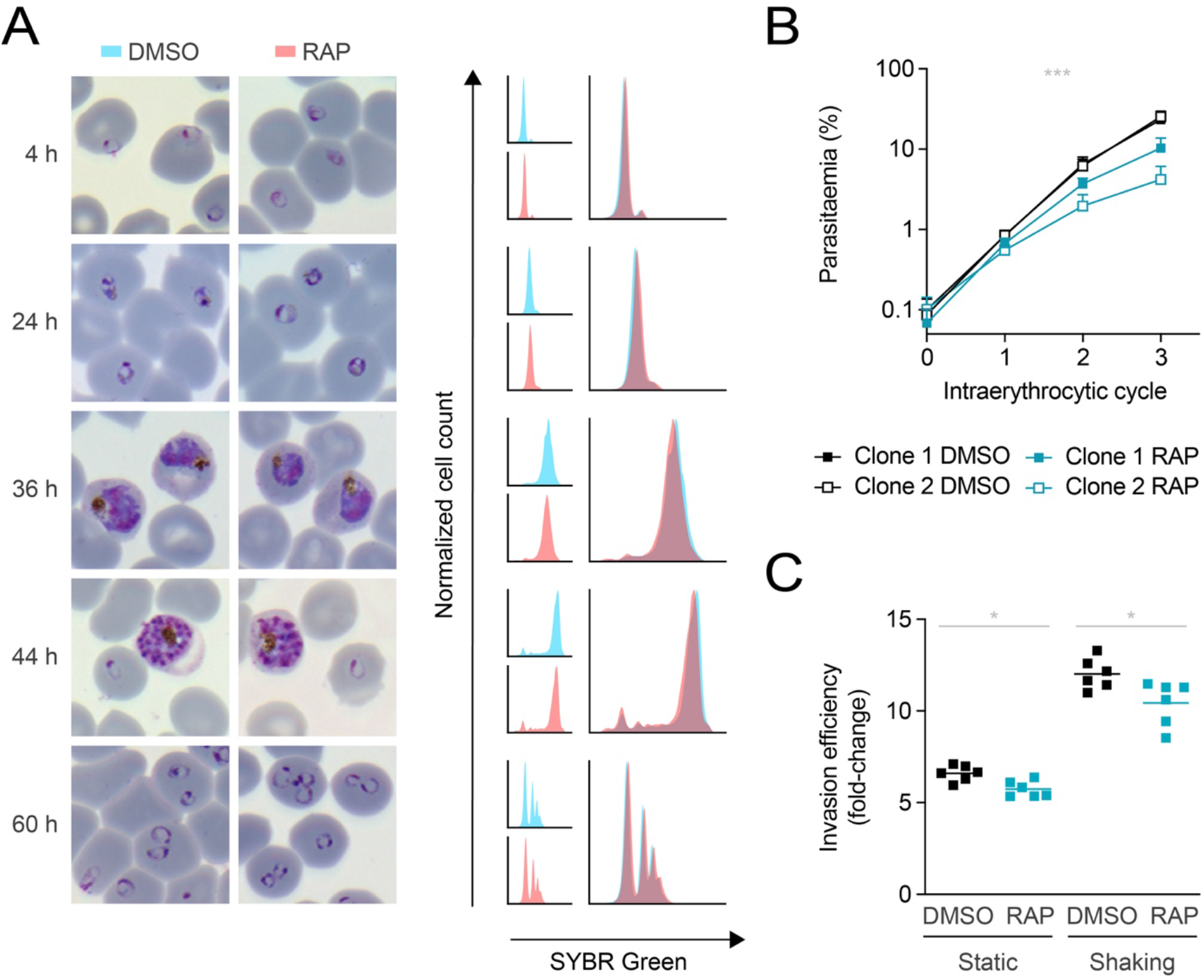
Impaired fitness of *in vitro* cultivated *Plasmodium falciparum* in the absence of PV5. (*A*) *Pf*PV5-deficient parasites mature normally *in vitro*. Tightly synchronized *pv5-3xHA:loxP* ring stages were treated with dimethyl sulfoxide (DMSO, blue) or rapamycin (RAP, red) and visualized by Giemsa staining 4, 24, 36, 44 and 60 hours later (left). In parallel, nuclear SYBR Green fluorescence was quantified by flow cytometry. Individual and merged histograms are depicted (right). Results are representative of two independent experiments. (*B*) Asexual parasite proliferation is impaired upon loss of *Pf*PV5. Shown are growth curves of two independent *pv5-3xHA:loxP* clonal lines upon treatment with DMSO or RAP, respectively. Averaged parasite multiplication rates are 7.6 (DMSO) and 4.6 (RAP). Shown are mean values +/-SD. ***, P<0.001; Two-way ANOVA. N=6 independent infections. (*C*) Impaired schizont to ring stage transition in the absence of *Pf*PV5. Schizonts from DMSO- and RAP-treated *pv5-3xHA:loxP* cultures were added to fresh erythrocytes and incubated under static or shaking conditions for 24 hours. Shown is the fold-change in parasitaemia, depicted as individual and mean values (bars). *, P<0.05; paired *t*-test. N=6 independent infections.

**Fig. S5.**
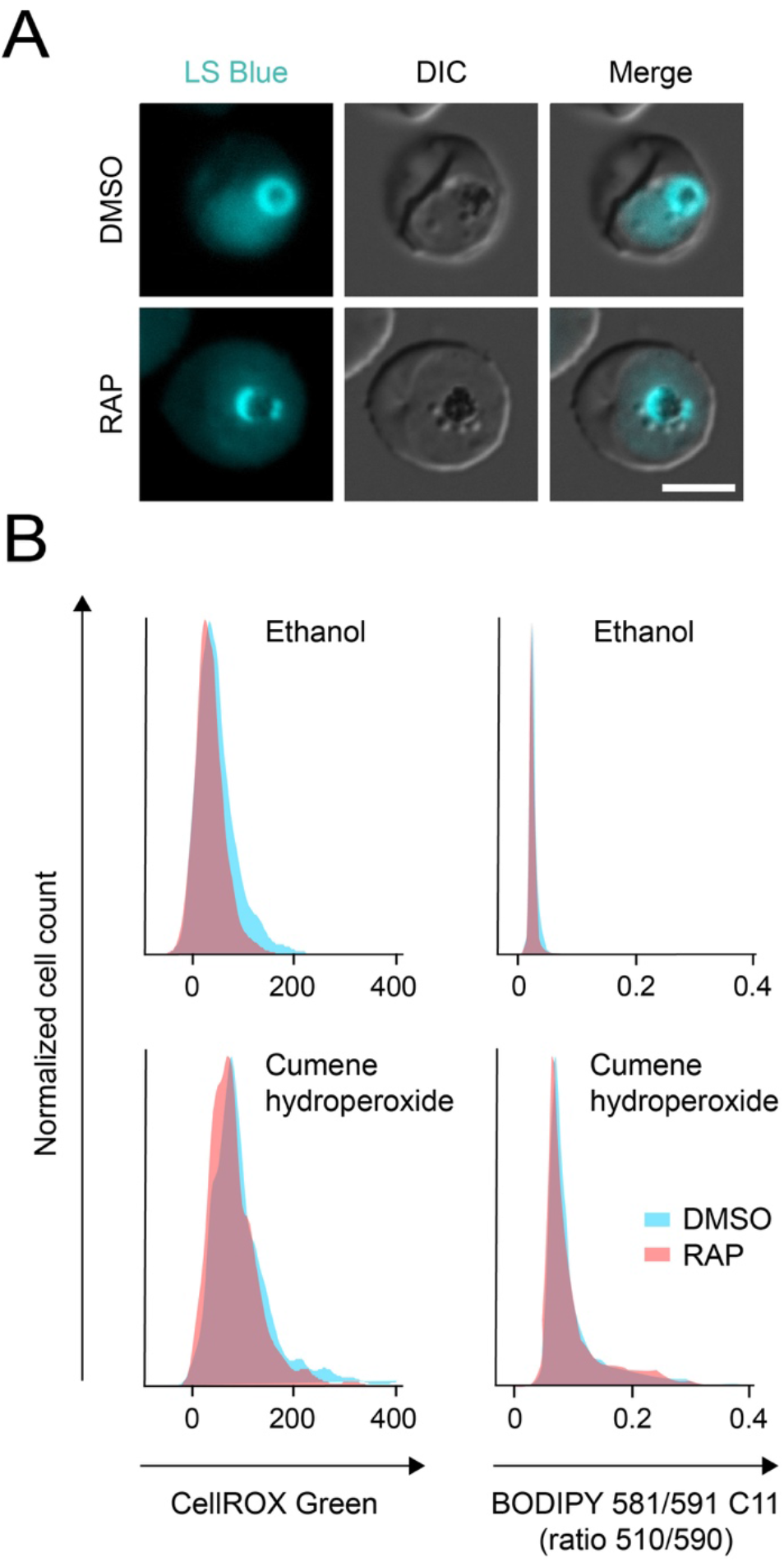
Absence of *Pf*PV5 does not cause dissipation of the vacuolar pH gradient nor an increase in oxidative stress. (*A*) *Pf*PV5-deficient parasites maintain an intact and acidic FV. Synchronized *pv5-3xHA:*loxP parasites were treated with dimethyl sulfoxide (DMSO) or rapamycin (RAP) from the ring stage onward and were stained with Lysosensor Blue DND-167 (LS Blue) 36 hours later. Shown are the LS Blue channel (cyan, left), differential interference contrast images (DIC, centre) and a merge (right). Bar, 5 μm. (*B*) No increased oxidative stress in the absence of *Pf*PV5. Synchronized *pv5-3xHA:loxP* parasites were treated with DMSO (blue) or RAP (red) from the ring stage onward, stained with the oxidative stress sensor CellROX Green (left) or with the ratiometric lipid peroxidation dye BODIPY 581/591 C11 (right) 36 hours later and analysed by flow cytometry. In addition, parasites had been treated with the oxidative stress-inducing agent cumene hydroperoxide (bottom) or with ethanol as the solvent control (top). Shown are the histograms of CellROX Green fluorescence intensity or of the 510/590 nm fluorescence ratio of BODIPY 581/591 C11. Results are representative of two independent experiments.

**Movie S1.** Absence of *Pf*PV5 ablates haemozoin movement within the food vacuole of *Plasmodium falciparum*.

Shown are differential interference contrast recordings of dimethyl sulfoxide (DMSO, left) and rapamycin-treated (RAP, right) *pv5-3xHA:loxP* parasites 36 hours following invasion. The video contains 120 frames shown at a 4x acceleration. Elapsed time is indicated in the upper right corner. Bar, 5 μm.

**Table S1.** Haemozoin crystal morphometry.

|  | WT | *pv5-3xHA:loxP* |  | WT vs.<br>DMSO | WT vs.<br>RAP | DMSO<br>vs. RAP |
| --- | --- | --- | --- | --- | --- | --- |
|  |  | DMSO | RAP |  |  |  |
| Area ($\mu\text{m}^2$) | 0.164 ($\pm 0.088$) | 0.107 ($\pm 0.052$) | 0.102 ($\pm 0.039$) | *** | *** | n.s. |
| Aspect ratio | 3.673 ($\pm 1.047$) | 2.781 ($\pm 1.061$) | 1.604 ($\pm 0.337$) | *** | *** | *** |
| Branching (%) | 0 | 27.5 | 96.1 |  |  |  |

n.s., non-significant; \*\*\*, $P < 0.001$ ; One-way ANOVA and Tukey's multiple comparison test. N>300 crystals.

**Table S2.**
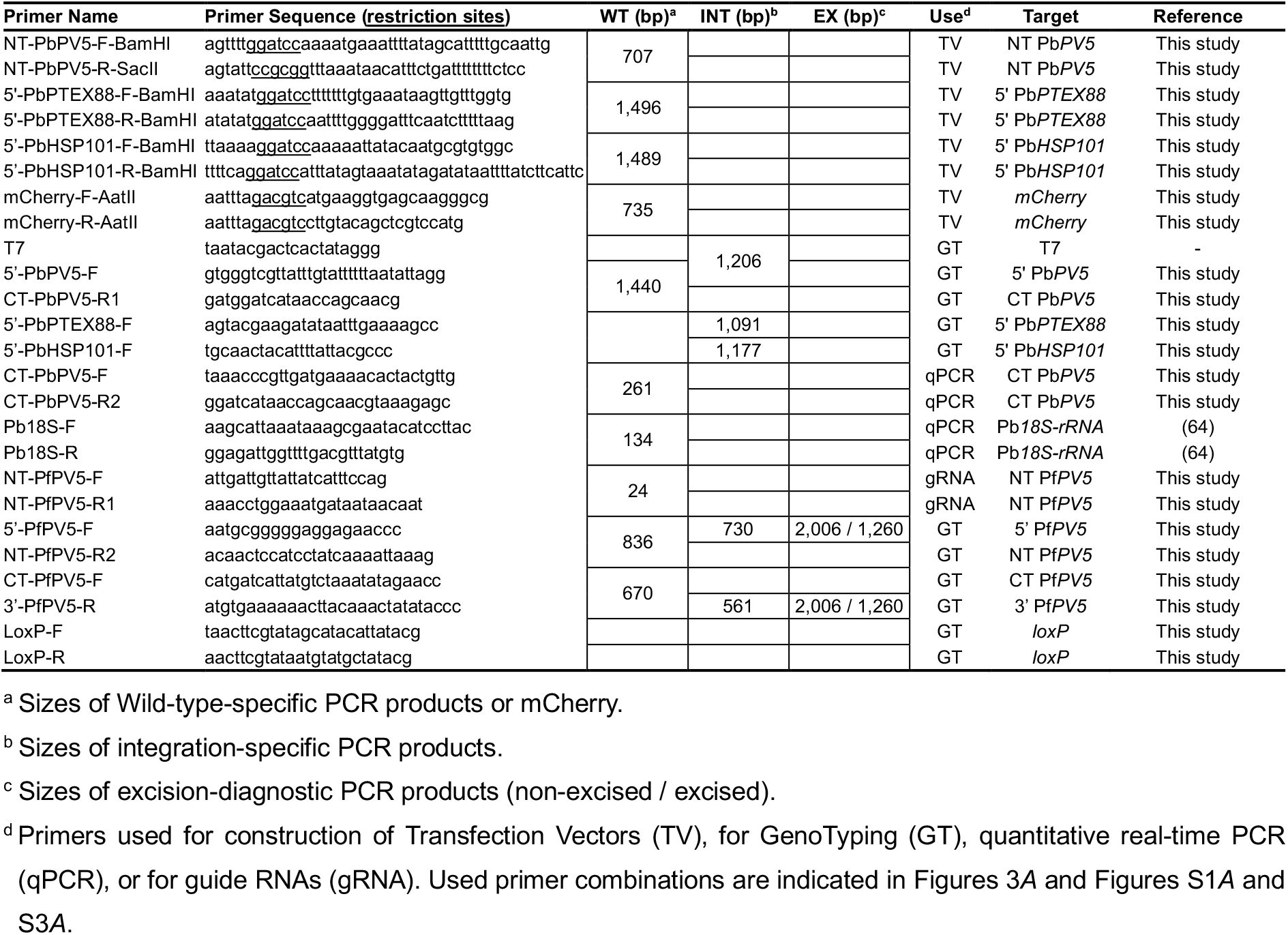
Primer sequences.

